# Orphan nuclear receptors Err2 and 3 promote a feature-specific terminal differentiation program underlying gamma motor neuron function and proprioceptive movement control

**DOI:** 10.1101/2022.01.25.477566

**Authors:** Mudassar N. Khan, Pitchaiah Cherukuri, Francesco Negro, Ashish Rajput, Piotr Fabrowski, Vikas Bansal, Camille Lancelin, Tsung-I Lee, Yehan Bian, William P. Mayer, Turgay Akay, Daniel Müller, Stefan Bonn, Dario Farina, Till Marquardt

**Author notes:** Co-correspondence. email id.

## Abstract

Motor neurons are commonly thought of as mere relays between the central nervous system and the movement apparatus, yet, in mammals about one-third of them function exclusively as regulators of muscle proprioception. How these gamma motor neurons acquire properties to function differently from the muscle force-producing alpha motor neurons remains unclear. Here, we found that upon selective loss of the orphan nuclear receptors Err2 and Err3 (Err2/3) in mice, gamma motor neurons acquire characteristic structural (e.g. synaptic wiring), but not functional (e.g. physiological firing rates) properties necessary for regulating muscle proprioception, thus disrupting gait and precision movements *in vivo*. Moreover, Err2/3 operate via transcriptional activation of neural activity modulators, one of which (Kcna10) promoted gamma motor neuron functional properties. Our work identifies a long-sought mechanism specifying gamma motor neuron properties necessary for proprioceptive movement control, which implies a ‘feature-specific’ terminal differentiation program implementing neuron subtype-specific functional but not structural properties.

**Summary:** The transcription factors Err2 and 3 promote functional properties in a subset of motor neurons necessary for executing precise movements.

## Introduction

Motor neurons tend to be thought of as passive links between the central nervous system and the movement apparatus by faithfully relaying descending neural commands to the skeletal musculature, where these signals evoke muscle contractions and thereby movement^1^. However, in zebrafish, motor neurons have been shown to provide active and direct feedback control for the neural networks driving rhythmic (swimming) locomotion^2^. In mammals, moreover, up to a third of the motor neurons do not connect to the force-generating extrafusal skeletal muscle fibers and do not directly contribute to muscle contractions^3-5^. Rather, these neurons, called gamma motor neurons, effectively contribute to the ability of mammals to learn and execute precision movements^6, 7^. Gamma motor neurons do so by regulating the flow of proprioceptive information the nervous system receives about skeletal muscle (and thus limb and trunk) position and velocity^8-11^ via direct synaptic connections onto the intrafusal fibers of stretch-sensitive mechanosensory organs called muscle spindles^3-6^. These motor neurons thereby facilitate ‘real-time’ adaptations of movements to changing external conditions by enhancing detection of discrepancies from intended movement trajectories^6, 7, 12^. In addition, gamma motor neurons generate muscle tone by recruiting the force-generating (alpha) motor neurons through a monosynaptic spindle afferent feedback ‘servo’ loop, thus assisting movement initiation and postural control^6^.

The ability of gamma motor neurons to control muscle proprioception entails their acquisition of intrinsic functional properties distinguishing them from the muscle force-generating alpha motor neurons^13^. For instance, their low firing thresholds and ability to rapidly gear up high firing rates appear to be exquisitely suited for achieving near-instant intrafusal fiber tension for maintaining or modulating muscle spindle dynamic range^6, 13^. Another feature allowing gamma motor neurons to effectively control muscle proprioception is their lack of monosynaptic (Ia) afferent feedback, which uncouples them from potential ‘short circuits’ by their own actions on muscle spindle activity^14, 15^. The acquisition of such specific combinations of functional (activation threshold, firing rate etc.) and structural properties (soma size, synaptic wiring preference etc.) inherent to neuron type or subtype identity^16^ is thought to involve overarching terminal differentiation (‘selector’) gene expression programs^17, 18^. Nevertheless, terminal differentiation programs have also been shown to promote only some but not all neuron subtype-specific structural features, like axonal versus dendritic wiring specificity^19^, raising the possibility that at least in some instances neuron type or subtype-specific properties could be specified by such programs in a modular (‘feature-specific’) manner.

Apart from their overall distinction from the gamma motor neurons, alpha motor neurons themselves exhibit a range of functional properties important for the adjustment of muscle force^20^. We and others have previously reported mechanisms underlying the diversification of alpha motor neurons proper, which involved both cell-autonomous actions by the non-canonical Notch ligand Dlk1^21^, as well as non-cell autonomous signals by region-specific astrocytes^22^. Yet, how gamma motor neurons acquire a unique suite of properties that allow them to control muscle proprioception, instead of generating muscle force, remained unaddressed. Here, we studied the specification of gamma versus alpha motor neurons in mice and identify the orphan nuclear receptors Err2 and Err3 as drivers of a ‘feature-specific’ terminal differentiation program underlying the acquisition of gamma motor neuron-specific functional but not structural properties required for proprioceptive movement control.

## Results

### Electrophysiological characterization of murine gamma motor neurons

Since the last recordings from mature gamma motor neurons dated from 1978 in the cat^13^, we first sought to establish a baseline for gamma motor neuron functional properties in the mouse. We were aided in this by a fortuitous finding that allowed us to selectively record gamma motor neurons in the mouse spinal cord based on the relative efficacy of retrograde Fluoro-Gold uptake (Figures 1A, 1B and S5A-S5G). We observed that indeed gamma motor neurons can be distinguished from alpha motor neurons based on their FG uptake as FG^high^ lacked sensory neuron innervations and expressed low or negligible levels of NeuN when compared to FG^low^ motor neurons (Figure 1B). Upon performing whole cell patch-clamp recordings at 20-22 days postnatally, we observed that the small-soma motor neurons with high-levels of Fluoro-Gold retention (FG^high^) exhibited an electrophysiological signature matching that previously reported for cat gamma motor neurons^13^ (Figures 1C-1E). The FG^high^ motor neurons showed a distinctive combination of lower rheobases with higher firing frequencies and gains, as well as higher instantaneous and steady-state firing rates when compared to the larger (FG^low^) motor neuron subtypes (Figures 1D, 1E and S6A and S6B and Supplementary Table S1). In contrast, FG^low^ motor neuron subtypes were characterized by a combination of low rheobases with low firing rates (alpha or beta motor neurons)^20^ (Figures 1D, 1E and S6A and S6B and Supplementary Table S1). Moreover, FG^high^ motor neurons showed significantly lower membrane input resistance, higher membrane capacitance and lower AHP-decay times when compared to FG^low^ motor neurons (Figures 1E and S6C and Supplementary Table S1). In addition to the overall quantitative differences in functional properties, gamma motor neurons in mouse as in the cat^13, 20^ further showed a distinctive combination of functional properties (e.g.: relatively very low activation thresholds + very high firing rates), compared to those of the slow (low thresholds + low firing rates) or fast alpha motor neuron subtypes (high thresholds + high firing rates).

**Figure 1.**
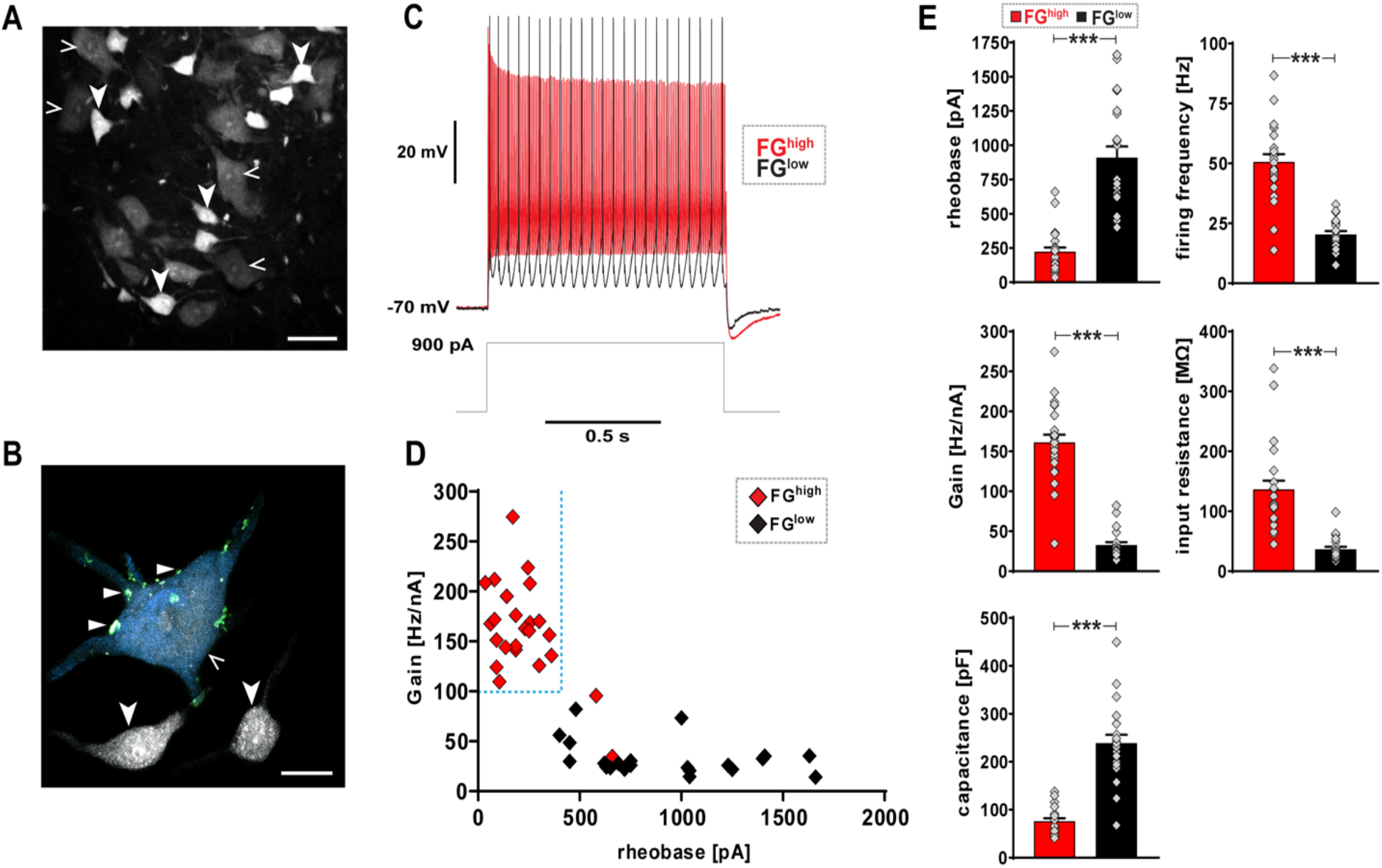
Direct electrophysiological interrogation of alpha and gamma motor neurons shows unique electrophysiological properties **(A-D)**. **(A)** Transversal section of P21 (wild-type) mouse lumbar spinal cord ventral horn: motor neurons labeled by retrograde tracer FluoroGold (FG). Putative alpha motor neurons retain low-levels of FG in soma (FG^low^) (open arrowheads), while putative gamma motor neurons retain high-levels of FG in soma (FG^high^) (arrowheads) (scale bar: 50 μm). **(B)** *Imaris* 3D reconstruction: vGlut1^+^ synaptic varicosities (triangles) associated with FG^low^ and NeuN^high^ alpha motor neuron (open arrowheads), but not with adjacent FG^high^ and NeuN^low or negligible^ gamma motor neurons (arrowheads) (scale bar: 20 μm). **(C)** Whole cell patch-clamp recordings: example traces of FG^high^ (black) and FG^low^ (gray) motor neurons upon 900 pA, 1 s square current pulse. **(D)** Scatter plot: FG^high^ (n=24, N=16) and FG^low^ (n=22, N=12) motor neurons exhibit divergent gamma and alpha subtype-defining electrophysiological signatures, respectively, including gamma subtype-specific combination with low rheobase and high gain by FG^high^ motor neurons (see Supplementary Table S1 for details). **(E)** FG^high^ motor neurons have significantly lower rheobase (pA) (221.87 ± 31.34), higher firing frequency (Hz) (50.65 ± 3.23), higher gain (Hz/nA) (161.01 ± 9.77), higher input resistance (136.62 ± 14.63) and lower capacitance (76.07 ± 6.01) when compared to FG^low^ motor neuron rheobase (909.09 ± 82.11), firing frequency (20.41 ± 1.70), gain (32.64 ± 4.02), input resistance (36.55 ± 4.16) and capacitance (239.1 ± 17.17), respectively (see Supplementary Table S1 for details). Data is presented as mean ± SEM. n= # of neurons, N= # of mice. Statistically significant differences between FG^high^ and FG^low^ neurons are indicated as: ***p<0.001, Student’s t-test).

### Correlated expression of orphan nuclear receptors Err2 and Err3 by gamma motor neurons

It had previously been established that the survival of gamma motor neurons beyond the early postnatal period relies on signals released by muscle spindles^23, 24^, which, in addition to the discovery of markers allowing *in situ* detection of gamma motor neurons^23, 24^ provided us with entry points for studying mechanisms underlying the diversification of motor neurons into alpha and gamma subtypes. We focused on the estrogen-related receptor (Err) subfamily of orphan nuclear receptors, which primarily function as ligand-independent transcription factors^25^, because of their contribution to cell type-specific functional (i.e.: electrophysiological) properties in other contexts^25^ and because of the previously reported expression of Err3 by gamma motor neurons^24^. We further found that the closely related Err3 paralogue Err2, with which it shares virtually the same DNA binding sequences^25^, was co-expressed with Err3 by gamma motor neurons (Figures 2A-2J and S2A-S2O) upon immunodetection using antibodies specifically recognizing either Err2 or Err3 (Figures S1A-S1Q), a co-expression which had been independently observed by others using single-cell RNAseq^26^. Through deeper analysis by quantitative immunodetection we indeed found high levels of correlated expression (Pearson correlation, r=0.86) of both Err2 and Err3 by motor neurons with relatively small somas characteristic for gamma motor neurons^27^ and low or negligible levels of the alpha motor neuron marker NeuN^23, 24^ (NeuN^low or negligible^) (Figures 2A-2J and S2A-S2O). The small-soma Err2/3^high^ NeuN^low or negligible^ motor neurons lacked vGlut+ varicosities on somatic or dendritic membranes (Figures 1B, 2L), indicating absence of monosynaptic spindle afferent input, a defining characteristic of gamma motor neurons^14, 15, 23, 24^. Similar to Err3, Err2 was initially broadly expressed by most motor neurons during embryonic development (Figures S3A-S3C), but high Err2 levels became increasingly confined to gamma motor neurons during the first two postnatal weeks (Figures S3D-S3R). Consistent with the dependency of gamma motor neuron maintenance on spindle-derived signals^23, 24^, adult motor neurons failed to retain high levels of both Err2/3 in Egr3-deficient mice with impaired muscle spindle development (Figures 2K, S3S-S3X and S3S’-S3X’). We noted that low Err2/3 levels were maintained by the remaining large-soma size NeuN^high^ motor neurons in Egr3-deficient mice (Figures S3S’-S3X’), while high Err2/3 levels persisted in a subset of ventral spinal interneurons in both Egr3-deficient mice (Figures S3S’-S3X’) and mice specifically lacking Err2/3 in motor neurons (Figure S1O-S1Q). Furthermore, *Imaris* reconstruction studies in control mice showed that indeed, FG^high^, Err2^high^, NeuN^low or negligible^ gamma motor neurons were contacted by few vGlut1^+^ terminals, while FG^low^, Err2^low or negligible^, NeuN^high^ alpha motor neuron somas were covered with vGlut1^+^ varicosities (Figures 2L-2N). Mouse gamma motor neurons thus exhibit high levels of correlated Err2 and Err3 expression.

**Figure 2.**
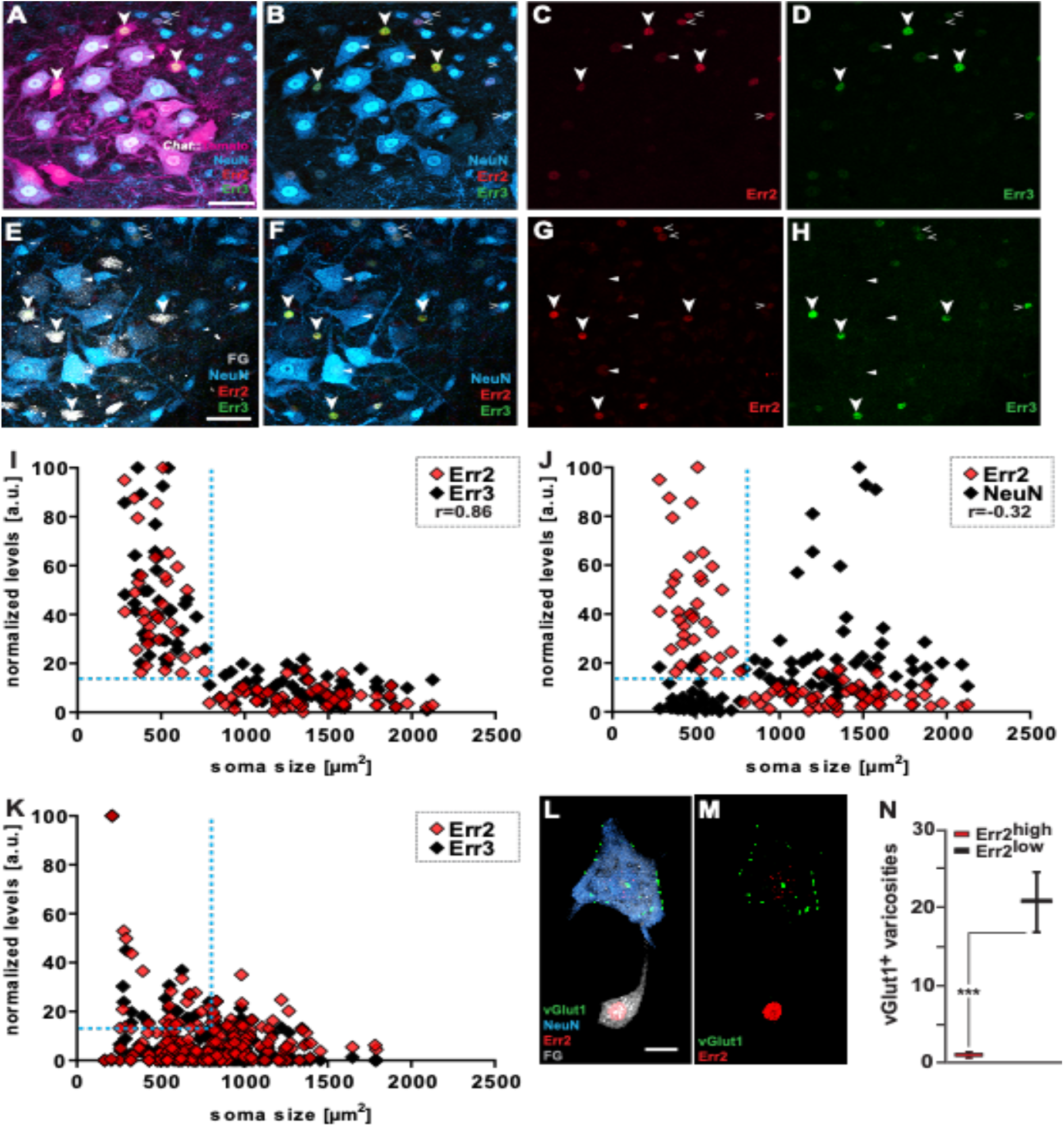
High levels of correlated Err2/3 expression by gamma motor neurons. **(A-D)**. Transversal section of P21 *Chat*^*Cre*^; *Rosa26*^*floxtdTomato*^ mouse lumbar spinal cord ventral horn: motor neurons genetically labeled by tdTomato (scale bar: 50 μm). Arrowheads: high Err2 and Err3 levels in small NeuN^low or negligible^, tdTomato^+^ motor neuron nuclei. Open arrowheads: relatively moderate-to-high-levels Err2/3 levels in NeuN^high^ tdTomato^-^ interneurons. Triangles: consistently lower but detectable levels in some large motor neurons with moderate-to-high NeuN levels. **(E-H)**. Transversal section of P21 (wild-type) mouse lumbar spinal cord ventral horn: motor neurons labeled by retrograde tracer FluoroGold (FG) (scale bar: 50 μm). Arrowheads: high Err2 and Err3 levels in small NeuN^low or negligible^ that retain high levels of FG^high^. Open arrowheads: relatively moderate-to-high levels Err2/3 levels in NeuN^high^ FG^-^ interneurons. Triangles: consistently lower but detectable levels in some large motor neurons with moderate-to-high NeuN levels. **(I)** Quantitative analysis: high Err2 (red) and Err3 (green) levels in motor neurons with small somas (n=93, N=3, Pearson’s correlation (r=0.86)). **(J)** High Err2 (red) and low NeuN (blue) levels in motor neurons with small somas (n=93, N=3, Pearson’s correlation (r=-0.32)). **(K)** Loss of high Err2 (red) and Err3 (green) levels by Egr3-deficient small motor neurons (n=184, N=3). **L-N)** Imaris 3D reconstruction: vGlut1^+^ synaptic varicosities associated with Err2^low or negligible^ NeuN^high^ motor neuron, but not with adjacent Err2^high^ NeuN^low or negligible^ motor neuron Err2^low^ motor neuron (motor neurons retrogradely labeled by FluoroGold, FG) (scale bar: 20 μm). **N)** Quantification of vGlut1^+^ synaptic varicosities associated with Err2^high^ or Err2^low^ motor neurons. N= # of mice and n= # of neurons. Statistically significant differences between motor neurons are indicated as: *p<0.05, **p<0.01, ***p<0.001, n.s.= not significant, Student’s t-test).

### Err2/3 are required for the acquisition of gamma motor neuron electrophysiological properties

Due to their correlated expression as well as their molecular similarity, we next asked whether Err2/3 would contribute to gamma versus alpha motor neuron functional diversification by selectively inactivating both *Esrrb* and *Esrrg* genes in motor neurons via Cre-mediated recombination in cholinergic neurons in *Esrrb*^*flox/flox*^; *Esrrg*^*flox/flox*^; *Chat*^*Cre*^ (Err2/3^cko^) mice (Figures S1O-S1Q). Because *Chat*^*Cre*^-mediated recombinase activity overlapped with endogenous Err2/3 expression in motor neurons but not in other cholinergic neuron types throughout the nervous system (Figures S4A-S4D), we concluded that Err2/3^cko^ mice permitted addressing Err2/3 function in motor neurons. Whole cell patch-clamp recordings performed in 20-22 days-old Err2/3^cko^ mice showed that most FG^high^ motor neurons failed to acquire a gamma motor neuron electrophysiological signature and instead shifted their properties dramatically towards resembling those of FG^low^ alpha motor neurons, with significantly elevated rheobases, as well as lowered gains and firing rates (Figures 3A, 3B and 3C, and S6D-S6H and Supplementary Table S1). At the same time, we did not observe significant changes in the properties of FG^low^ alpha motor neurons in Err2/3^cko^ mice when compared to control mice (Figures 3D, 3E and 3F and S6I-S6M and Supplementary Table S1). Err2/3 thus appear to be prerequisite for the acquisition of gamma motor neuron functional properties, but not for the development of morphologically distinguishable gamma motor neurons proper.

**Figure 3.**
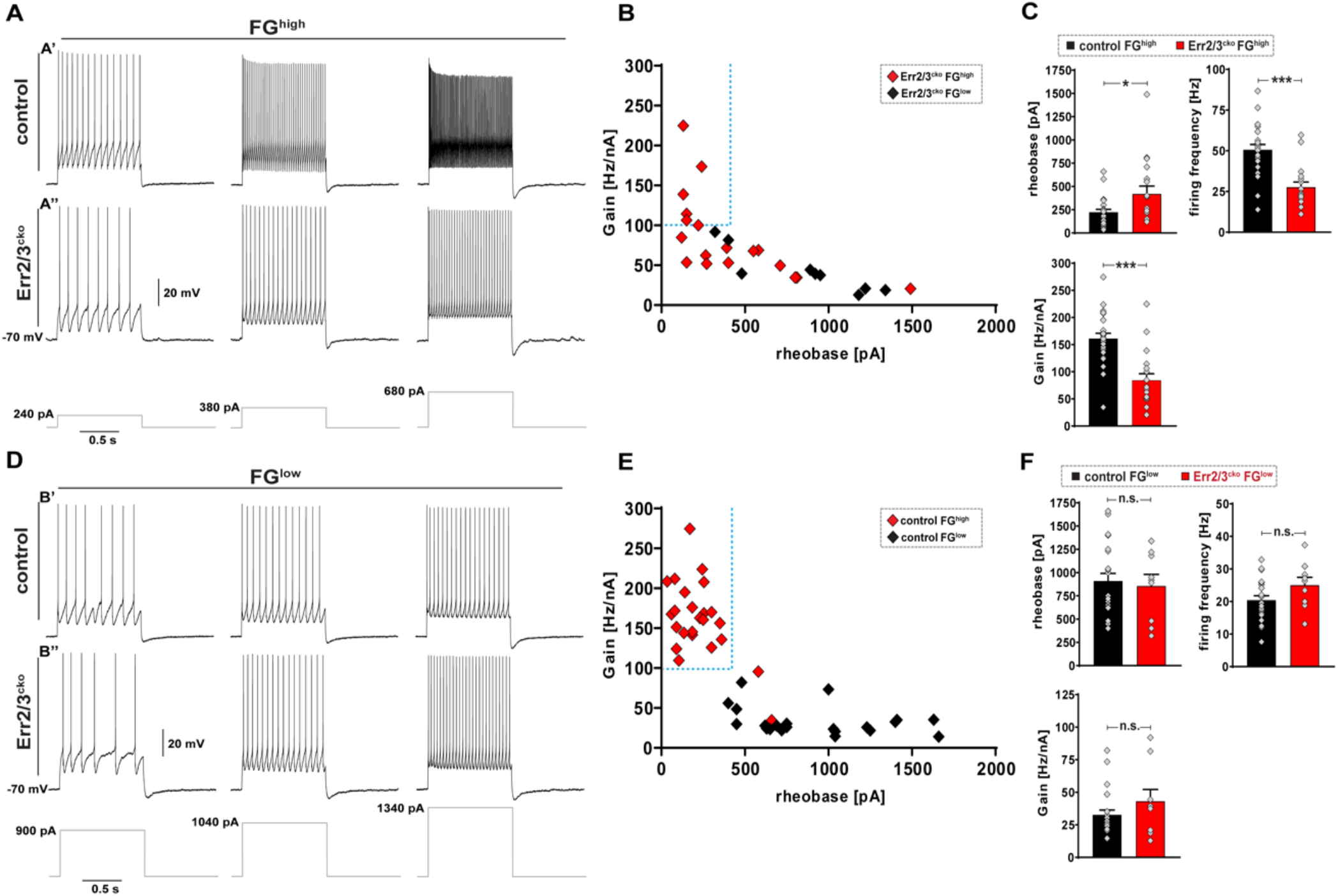
Err2/3 are required for the acquisition of a gamma motor neuron functional properties. **(A)** Example traces of whole cell patch-clamp recordings upon 240, 380 or 680 pA, 1 s square current pulses from small soma motor neurons exhibiting high-levels of FG incorporation (FG^high^) from control mice or Err2/3^cko^ mice. Control FG^high^ motor neurons **(A’)** exhibit higher firing rates compared to FG^high^ Err2/3^cko^ motor neurons **(A’’)** and gear up their firing rates more rapidly in response to current pulses. **(B)** Scatter plot: lack of segregation of Err2/3^cko^ FG^high^ gamma motor neuron and Err2/3^cko^ FG^low^ alpha motor neuron electrophysiological signatures. **(C)** Err2/3^cko^ FG^high^ motor neurons show higher rheobase (419.72 ± 84.11), lower firing frequency (27.61 ± 3.15) and lower gain (84.01 ± 12.34) compared control FG^high^ motor neuron rheobase (221.87 ± 31.34), firing frequency (50.65 ± 3.23), gain (161.01 ± 9.77), respectively. **(D)** Example traces of whole cell patch-clamp recordings upon 900, 1040 or 1340 pA, 1 s square current pulses from large soma motor neurons with lower levels of FG incorporation (FG^low^) from control mice or Err2/3^cko^ mice. Err2/3^cko^ FG^high^ motor neurons exhibit comparable firing rates and properties to control FG^low^ motor neurons. **(E)** Scatter plot: segregation of electrophysiological signatures between control FG^high^gamma motor neurons versus control FG^low^ alpha motor neurons. (Note: Data from is from Fig. 1D). **(F)** No significant differences seen in Err2/3^cko^ FG^low^ alpha motor neuron subtype rheobase (855.55 ± 124.85), firing frequency (25.03 ± 3.76) and gain (43.04 ± 9.81) when compared to control FG^low^ alpha motor neuron subtype rheobase (909.09 ± 82.11), firing frequency (20.41 ± 1.70), gain (32.64 ± 4.02), respectively (see Supplementary Table S1 for details). Data is presented as mean ± SEM. n= # of neurons and N= # of mice. Statistically significant differences between FG^high^ and FG^low^ neurons are indicated as: *p<0.05, **p<0.01, ***p<0.001, n.s.= not significant, Student’s t-test).

### Err2/3-dependent gamma motor neuron functional properties are required for movement accuracy

The biophysical properties of gamma motor neurons are thought to be exquisitely suited to effect instant intrafusal fiber peak tension for maintaining and modulating spindle dynamic range and thus muscle proprioception^13^. We therefore asked how a shift towards an alpha motor neuron-like electrophysiological signature in gamma motor neurons would impact movements relying on proprioceptive feedback from muscles. Because of the exclusive association of high Err2/3 levels with gamma motor neurons, and the lack of a significant impact of Err2/3 loss on other motor neuron subtypes, we predicted that beta motor neuron function would be preserved in Err2/3^cko^ mice, thus allowing us to study the contribution of gamma motor neuron function to movement control in the otherwise intact animal. Similar to Egr3-deficient mice^9, 10^, Err2/3^cko^ mice exhibited marked postural and gait alterations, including changes in metrics related to foot placement, weight bearing, stride, stance, braking and propulsion (Figures 4A, 4B, S8D and S8E), consistent with the predicted contributions of gamma motor neuron-assisted spindle function to posture, gait phase-transitions and force generation during locomotion^6, 9-11^. Err2/3^cko^ mice were nevertheless able to sustain the same range of speeds as control mice in a treadmill locomotion task with little dependency on muscle proprioception^9, 10^ (Figures S8A-S8C). However, Err2/3 loss from motor neurons triggered a failure to handle precision tasks with extensive reliance on muscle proprioception^9, 10^, such as navigating a narrow horizontal beam (Figures 4C, 4D and Supplementary Movies S1 and S2) or a horizontal ladder (Figures 4E, 4F and Supplementary Movies S3 and S4), indicating that the electrophysiological signature implemented by Err2/3 in gamma motor neurons is prerequisite for effective modulation of muscle proprioception, and thereby the execution of precision movements. To further test this idea, we recorded spindle afferent responses via suction electrodes in nerve-muscle preparations^31^ derived from Err2/3^cko^ or control mice. In these preparations muscle stretch applied by a force transducer elicited similar Ia afferent responses in Err2/3^cko^ and control mice (Figures 4G, 4G’ and S7A-S7D), consistent with the morphologically normal spindle assembly in these animals (Figures 5K and S7G, S7H). In contrast to control spindles (Figures S7A and S7B), however, Err2/3^cko^ spindle afferents frequently exhibited reduced firing rates at muscle resting length (Figures 4H, 4H’ and S7C, S7D), possibly due to a decrease in basal intrafusal fiber contractility caused by chronic disruption of gamma motor neuron input. The Err2/3-dependent implementation of gamma motor neuron functional properties therefore appears to be prerequisite for regulating spindle-mediated muscle proprioception and movement control.

**Figure 4.**
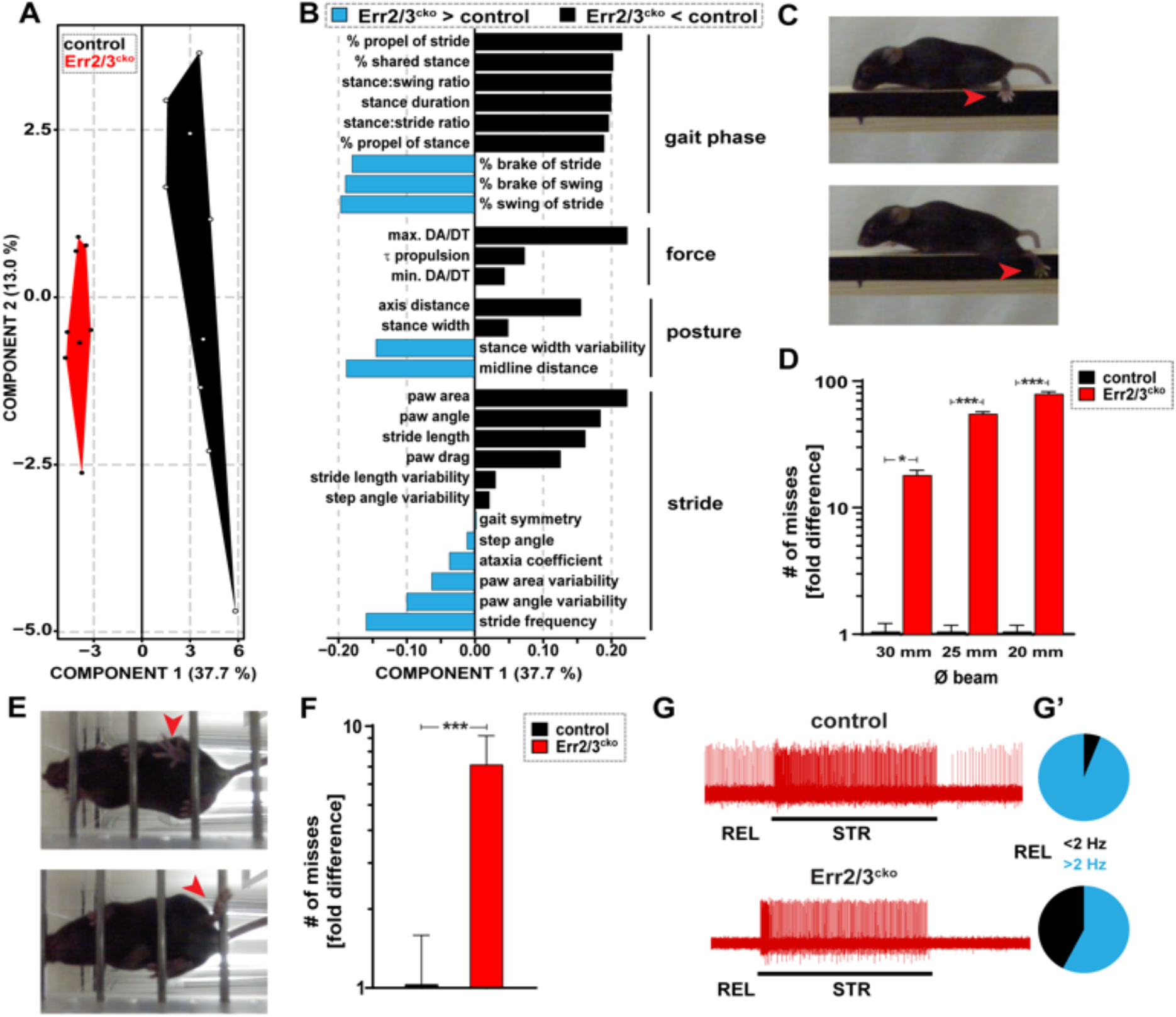
Err2/3 is required for the acquisition of gamma motor neuron-specific functional properties and the execution of precision movements. **(A, B) (A)** Polygon graphs based on partial least squares (PLS) analysis of 58 gait variables measured during treadmill locomotion at 25 m•s^-1^ reveals significant gait alterations in Err2/3^cko^ mice (n=8, 3 trials/animal) when compared to control mice (n=9, 3 trials/animal). Each dot represents a single animal, polygons group animals of the same genotype, the segregation of which along the x-axis indicate that Err2/3^cko^ mice exhibit significant gait alterations at all speeds tested (Err2/3^cko^ mice, 3 trials each). **(B)** Negative (black) or positive (light blue) changes (arbitrary units) in gait variables in Err2/3^cko^ compared to control mice ranked by predictive value independent of sign, including reduced stance-swing phase ratio (gait phase related), reduced propulsion velocities (force related), decreased stance width (posture related), increased paw angle variability (stride related). **(C)** Still images of Err2/3^cko^ mouse navigating a horizontal beam. Red arrow: (example of a “miss”): foot missing the beam during swing-stance transition, causing the hindlimb and animal to slip during swing phase (red arrow in **C**). **(D)** Err2/3^cko^ (18.0 ± 1.75, 55.0 ± 2.20, 79.0 ± 3.49) but not control (1.0 ± 0.175, 1.0 ± 0.125, 1.0 ± 0.125) mice exhibit dramatically increasing erratic locomotion (18, 55 and 79-fold increase in the # of misses) upon navigating horizontal beams with decreasing width 30 mm, 25 mm and 20 mm, respectively (control N=4, Err2/3^cko^ N=4, 4-5 trials/animal). **(E)** Still images of Err2/3^cko^ mouse navigating a horizontal ladder. Examples of a “miss” (red arrow in upper panel): foot missing a rung during swing-stance transition, causing the hindlimb to slip during swing phase. Note: animals frequently attempted to compensate such misses by using the “slipped” hindlimb to push against the rung and propel itself forward (red arrow in lower panel). **(F)** Err2/3^cko^ (7.12 ± 2.06) exhibit significantly more erratic locomotion (7-fold increase in the # of misses) when compared to control (1.0 ± 0.55) upon navigating horizontal ladder (control N=5, Err2/3^cko^ N=5, 4-5 trials/animal). **(G)** Example traces showing Ia spindle afferent responses during resting length (REL) or stretch (STR) applied by force transducer: normal STR responses, but reduced REL firing of Ia afferents in Err2/3^cko^ mice (N=10) when compared to control mice (N=10). (**G’**) Pie charts: ratio of Ia afferents firing above or below 2 Hz at REL (total trials: control n=84, Err2/3^cko^ n=118; control >2 Hz n=79, Err2/3^cko^ >2 Hz n=68). Data is presented as mean ± SEM. N= # of mice, n= # of 1a afferents. Statistically significant differences between control and Err2/3^cko^ mice are indicated as: *p<0.05, **p<0.01, ***p<0.001, n.s.= not significant, Student’s t-test).

**Figure 5.**
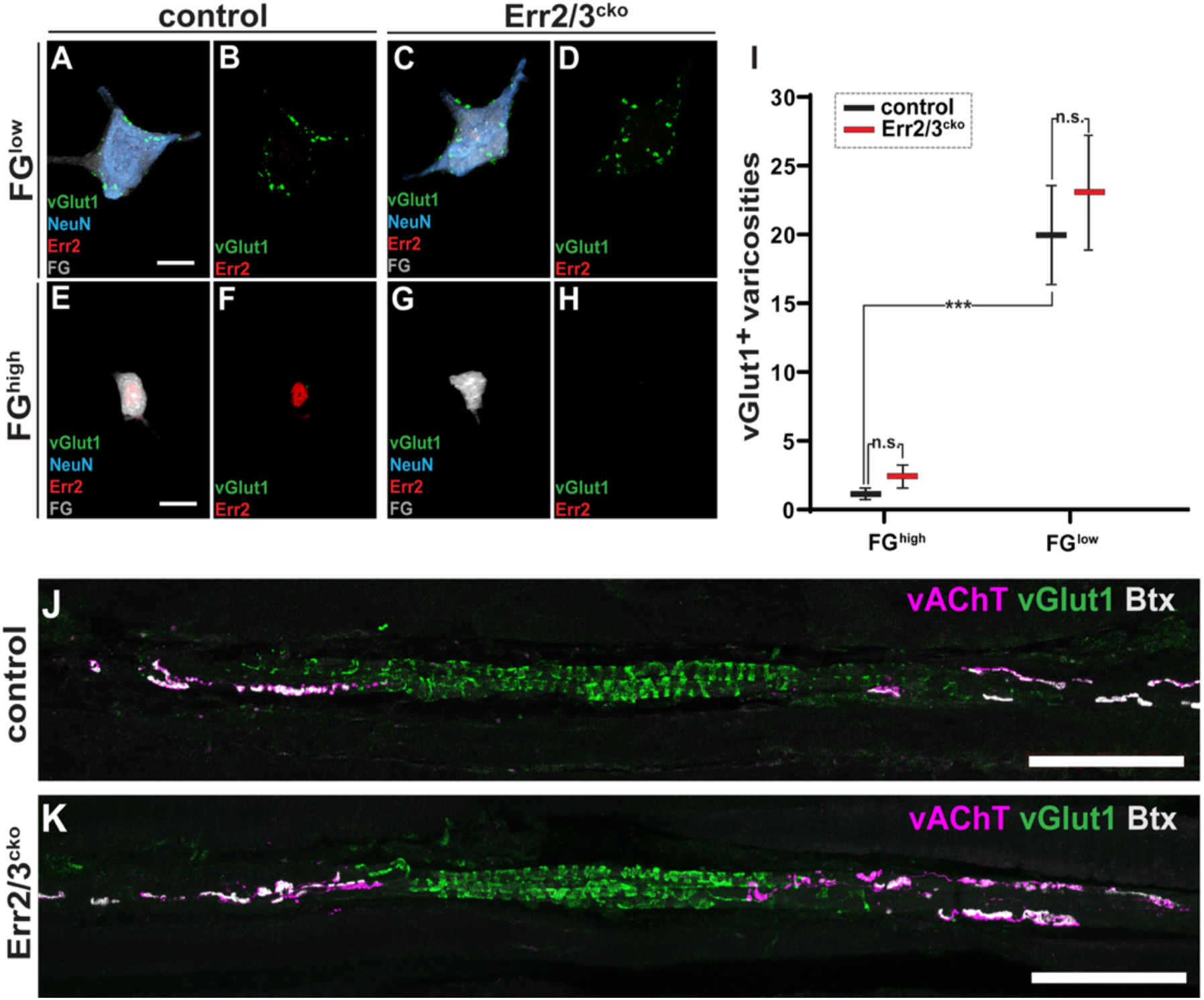
Acquisition of gamma motor neuron-specific morphology and synaptic connectivity without Err2/3. **(A-I)** Imaris 3D reconstruction: vGlut1^+^ synaptic varicosities (green) associated with large FG^low^, NeuN^high^ motor neurons in control **(A, B)** and Err2/3^cko^ mice **(C, D)** (scale bar: 20 μm). **(E-H)** Absence of vGlut1^+^ puncta on FG^high^, NeuN^low or negligible^ motor neurons in control **(E, F)** and Err2/3^cko^ mice **(G, H)** (scale bar: 20 μm). **(I)** Significant difference in vGlut1^+^ varicosities associated with FG^high^, NeuN^low or negligible^ and FG^low^, NeuN^high^ motor neurons from control mice. Lack of significant differences in vGlut1^+^ synaptic varicosities associated FG^low^, NeuN^high^ motor neurons in control (20 ± 3.48, n=9) versus Err2/3^cko^ mice (23.1 ± 4.06, n=10). Lack of significant differences in vGlut1^+^ varicosities between control FG^high^, NeuN^low or negligible^ (1.1 ± 0.36, n=10) and Err2/3^cko^ FG^high^, NeuN^low or negligible^ motor neurons (2.4 ± 0.78, n=10). **(J, K)** P70 mouse extensor digitorum longus (EDL) muscle spindles of control **(J)** and Err2/3^cko^ **(K)** mice. **(J)** Distribution of Ia sensory annulospiral endings in the central spindle segment (visualized by vGlut1, green), motor innervation (vAChT, magenta) and their postsynaptic sites (Btx, alpha bungarotoxin, grey) (scale bar: 100 μm). **(K)** Normal appearance of Err2/3^cko^ muscle spindle based on the distribution of sensory and motor innervation (scale bar: 100 μm). Data is presented as mean ± SEM. n= # of neurons. Statistically significant differences control and Err2/3^cko^ motor neurons are indicated as: *p<0.05, **p<0.01, ***p<0.001, n.s.= not significant, Student’s t-test).

### Lack of Err2/3 in gamma motor neurons does not alter their development and connectivity patterns

Since we observed that the loss of Err2/3 in gamma motor neurons in Err2/3^cko^ mice led to the loss of a gamma motor neuron electrophysiological identity, including low rheobases, high firing frequencies and gains, we then asked whether Err2/3 are necessary for gamma motor neuron morphology and connectivity. In addition to electrophysiological features, gamma motor neurons characteristically possess small soma sizes, lack 1a sensory neuron pre-synaptic innervation and send axons that innervate muscle spindle intrafusal fibers^24,27^. Remarkably, the lack of Err2/3 in Err2/3^cko^ mice did not affect the development of small-soma FG^high^ and NeuN^low or negligible^ motor neurons (Figures S3D-S3R). Similar to control mice (Figures 5E, 5F and 5I), the small-soma FG^high^, Err2^-^, NeuN^low or negligible^ motor neurons in Err2/3^cko^ mice (Figures 5G, 5H and 5I) lacked vGlut1^+^ varicosities on somatic or dendritic membranes. Moreover, FG^low^, Err2^-^, NeuN^high^ motor neurons retained their vGlut1^+^ terminals in both Err2/3^cko^ mice (Figures 5C, 5D and 5I) when compared to control mice (Figures 5A, 5B and 5I). We next observed the morphology of the muscle spindles. The muscle spindle appearance, notably the innervation by motor axon presynaptic termini within the peripheral contractile segments of the intrafusal muscle fibers was preserved in Err2/3^cko^ muscle spindles (Figures 5K) when compared to control mice (Figures 5J). Taken together, we surmise that Err2/3 are not necessary for gamma motor neuron structural identity, since the lack of Err2/3 in gamma motor neurons did not affect the development of morphologically distinct gamma motor neurons and did not perturb gamma motor neuron pre- and post-synaptic innervation patterns in Err2/3^cko^ mice.

### Err2/3 are sufficient in forcing gamma motor neuron functional properties in chick

We next tested whether Err2/3 would also be sufficient to promote a gamma motor neuron electrophysiological signature by performing whole cell patch-clamp recordings on chick motor neurons^21^ engineered to stably express elevated Err2/3 levels (Figures 6A-6D’). Indeed, forced expression of Err2/3 partially shifted motor neuron properties towards an electrophysiological signature recapitulating that of mouse or cat gamma motor neurons^13,20^, including a combination of high gains and firing rates (Figures 6E-6I, S6N and Supplementary Table S2), suggesting that high Err2/3 levels are not only necessary but also sufficient to promote a gamma motor neuron electrophysiological signature. This effect on motor neuron properties was enhanced by fusing Err2 to the heterologous transcriptional activation domain VP16 (Figures 6G-6I, S6N and Supplementary Table S2), but not by its fusion to the *engrailed* transcriptional repressor domain (EnR) (Figures 6G-6I, S6N and Supplementary Table S2), suggesting that in this context Err2 primarily functions as a transcriptional activator. Thus, Err2/3 appear to be not only necessary but also sufficient to promote gamma motor neuron functional properties by operating as transcriptional activators.

**Figure 6.**
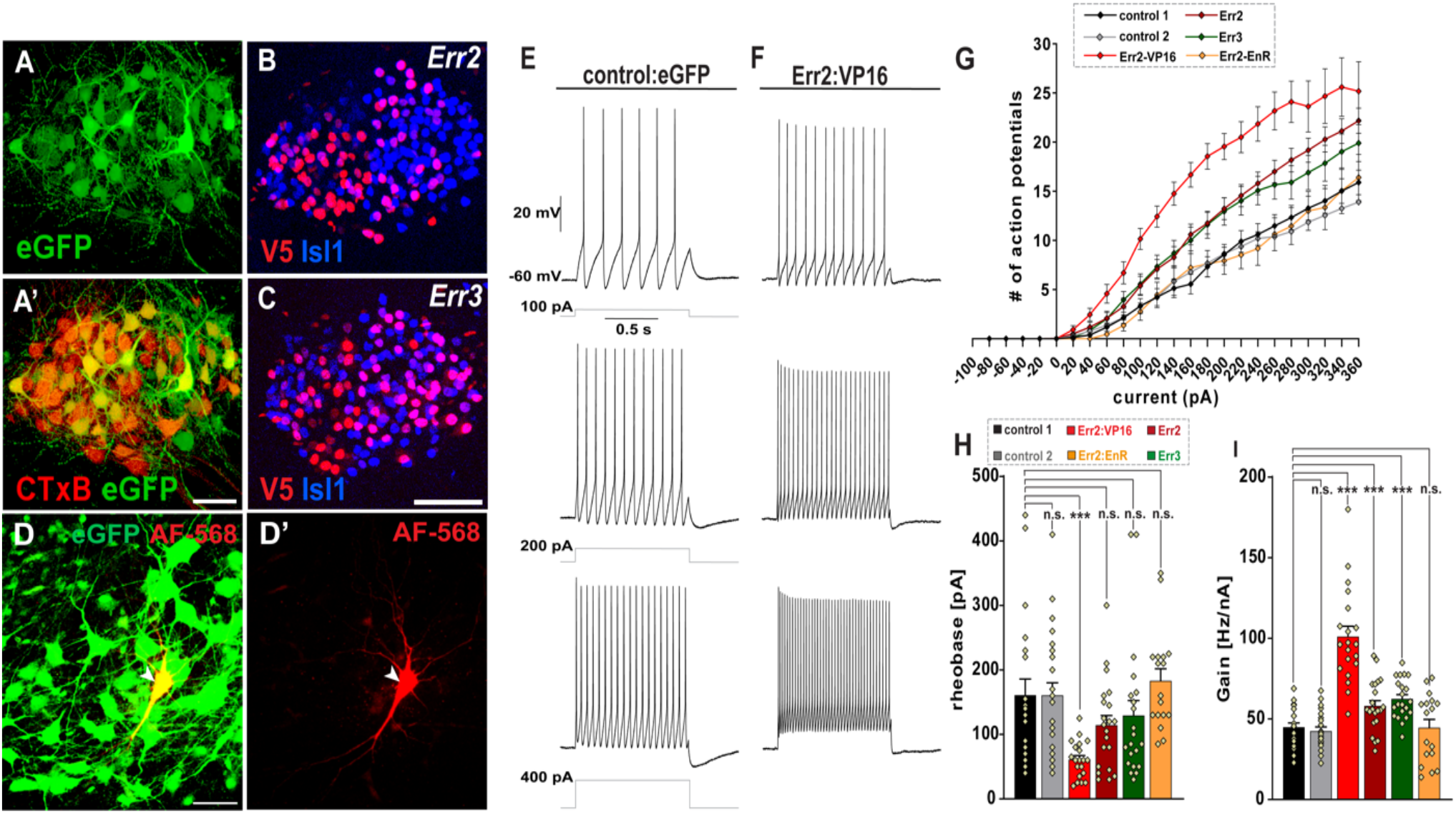
Err2/3 promote gamma motor neuron-specific functional properties through transcriptional activation. **(A-D’)** Overview of E12 chick spinal cord: stable transfection with expression vector driving eGFP expression **(A)** in ventral horn motor neurons. Higher magnification of eGFP expressing neurons further identified as motor neurons by retrograde cholera toxin B (CTxB) tracing upon *in ovo* injection into the hindlimb **(A’)** (scale bar: 100 μm). Examples of expression and nuclear localization of mouse Err2 **(B)** and Err3 **(C)** in chick motor neurons (detected via N-terminal V5 epitope tags) (scale bar: 30 μm). eGFP^+^ motor neuron recorded with patch pipette containing Alexa fluor-568 dye (red) **(D and D’)** (scale bar: 50 μm). **(E, F)** Example traces of current clamp recordings of chick motor neurons (in acute spinal cord slice preparations) forcedly expressing eGFP only (control) **(E)** or Err2:VP16 and eGFP **(F)**, upon 100, 200 and 400 pA, 1s square current pulses: motor neurons expressing elevated Err2 levels exhibit higher firing rates and gear up their firing rates more rapidly in response to current pulses **(F). (G)** Current-action potential response curves for control transfected motor neurons (black, n=21). Similar leftward shifts in the slope of the response curve upon expressing Err2 (dark red, n=21) or Err3 (green, n=21). Exaggerated leftward shift in the slope of the response curve upon expressing Err2 fused to a transcriptional activation domain (Err2:VP16) (red, n=20), while lack of shift when expressing Err2 fused to a transcriptional repression domain (Err2:EnR) (orange, n=17). **(H)** Compared to control 1 (160.95 ± 25.85), significant decrease in rheobase was observed upon over-expression of Err2:VP16 (61 ± 6.36), while no significant change in rheobase was observed upon over-expression of Err2 (114.28 ± 15.10), Err3 (129.28 ± 23.41) or Err2:EnR (182.94 ± 18.81). **(J)** Compared to control 1 (44.84 ± 2.75), significant increase in gain was observed upon over-expression of Err2:VP16 (101.01 ± 6.58), Err2 (58.00 ± 3.49), Err3 (62.47 ± 2.66) but not upon the forced expression of Err2:EnR (44.8 ± 5.14). No differences detected between control 1 and control 2 (n=with parameters recorded during two different experiments at different time points upon expressing “eGFP only” control vectors, thus demonstrating robustness of the assay. Data is presented as mean ±SEM. n= # of neurons. Statistically significant differences are indicated as: *p<0.05, **p<0.01, ***p<0.001, n.s.= not significant, Student’s t-test).

### Err2/3 drive gamma motor neuron functional properties by activating the expression of the shaker K^+^ channel subunit gene *kcna10*

Reasoning that in these gain-of-function experiments in chick, Err2 would likely operate through the same intermediate factors through which it would normally tune motor neuron electrophysiological properties, we performed comparative transcriptome profiling by RNA sequencing of chick motor neurons forcedly expressing Err2 (Figures 7A and S9G-S9J, Supplementary Table S3). In these experiments, elevated Err2 levels significantly activated a set of genes largely distinct from the gene signature activated by the previously identified (fast) alpha motor neuron determinant Dlk1^21^ (Figures 7A and S9G-S9J, Supplementary Table S3). This included activation by Err2 of *Kcna10*, which encodes a member of the shaker family of voltage-gated K^+^ channels implicated in neuronal excitability^29^ (Figures 7A and S9H), the promoter of which contained an evolutionary conserved region (ECR) with three clustered Err2/3 DNA binding motifs (Figures 7B and S9K). In chick motor neurons, moreover Err2 boosted reporter gene activity driven by the *Kcna10* ECR (Figures 7C, S9C and S9D), but not upon introducing mutations into the ECR’s Err2/3 binding motifs (Figures 7B, 7C, S9E and S9F). Ultimately, similar to *Err2*, forced expression of *Kcna10* shifted motor neuron electrophysiological properties towards lower rheobases and higher firing rates typical for gamma motor neurons (Figure 7D and Supplementary Table S2), together suggesting that Err2/3, like Dlk1^21^ or Islet in *Drosophila*^*30*^, chiefly operate through voltage-gated K^+^ channels to promote neuron subtype-specific electrophysiological properties (Figure 7E). Taken together, these data suggest that Err2/3 operate as transcriptional activators to implement a feature-specific gene expression program encoding neural activity modulators to promote gamma motor neuron functional but not structural properties.

**Figure 7.**
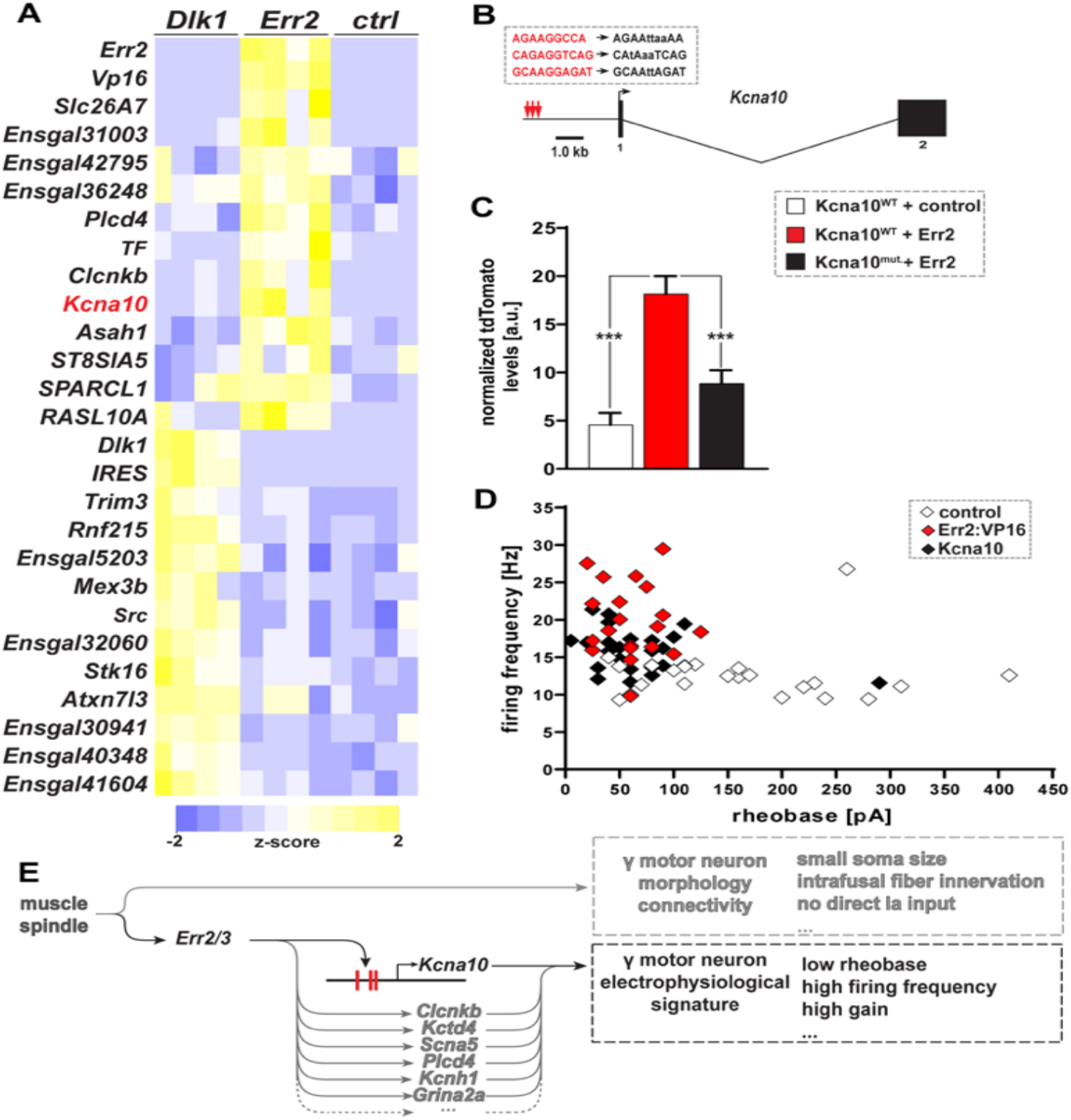
Err2 operates through *Kcna10* to promote a gamma motor neuron electrophysiological signature. **(A)** Heatmap based on transcript reads (transcript per million (TPM)) detected by RNA sequencing: Err2 (N=4) and Dlk1 (N=4) promote different gene expression signatures in chick motor neurons when compared to control (N=4), including *Kcna10* (red) by Err2 (only upregulated genes are shown). **(B)** *Kcna10* genomic locus: 3 clustered Err2/3 binding sites within promoter region. **(C)** Reporter tdTomato fluorescence driven by wild-type (WT) *Kcna10* promoter is boosted by Err2 co-transfection (18.14 ± 1.85) (red, n=102) in chick motor neurons when compared to control (4.57 ± 1.23) (gray, n=100) and decrease in tdTomato reporter fluorescence upon mutating the Err2/3 binding sites (8.85 ± 1.38) (black, n=101). **(D)** Whole cell patch-clamp recordings: forced Err2:VP16 expression (red, n=20) shifts chick motor neuron properties towards a gamma motor neuron-like electrophysiological signature (high firing rates, low rheobases) compared to control (white, n=23). Forced *Kcna10* (black, n=24) expression recapitulates the promotion of a gamma motor neuron electrophysiological signature by Err2 in chick motor neurons when compared to control (white, n=23) motor neurons. **(E)** Summary: Err2/3 bind 3 clustered binding sites within the *Kcna10* promoter region to drive *Kcna10* expression that promotes gamma motor neuron electrophysiological signature of low rheobases and high firing rates while not affecting the morphology and connectivity identities of gamma motor neurons. Data is presented as mean ± SEM. n= # of neurons, N= # of embryos. Statistically significant differences are indicated as: *p<0.05, **p<0.01, ***p<0.001, n.s.= not significant, Student’s t-test).

## Discussion

Neuronal specification is thought to involve gene expression programs that coordinate the acquisition of both functional (activation threshold, firing rate etc.) and structural properties (soma size, synaptic wiring preference etc.) associated with neuron type or subtype identities^16-18^. Here, we have shown that during the diversification of motor neurons into gamma and alpha subtypes, the acquisition of neuron subtype-specific functional properties can (at least to a large degree) be achieved independently from the acquisition of structural properties (Figure 7E). The disruption of proprioceptive movement control we observed upon eliminating gamma motor neuron-specific functional (but not structural) properties therefore suggests that homeostatic plasticity, which to some extent can compensate for quantitative fluctuations in neural activity^28^, is unable to offset a qualitative shift in the biophysical properties of one neuron subtype towards those of another. Since the ‘feature-specific’ action of Err2/3 in specifying neuron subtype-specific properties is apparently important for the function of the entire neural network involved in muscle spindle-dependent movement control, it will be interesting to determine whether a similar separation of functional and structural terminal differentiation programs exists elsewhere in the mammalian nervous system. This, in turn, would open the intriguing possibility of utilizing ‘feature-specific’ terminal differentiation factors as indicators for a certain set of biophysical properties (i.e.: as ‘markers for neuronal function’), instead of as markers for neuron type or subtype identities proper.

Two types of gamma motor neuron output, static or dynamic, are thought to normally tune muscle spindle sensitivity during different movement tasks^6, 33^. Three observations led us to conclude that the actions of Err2/3 do not distinguish between both types of gamma motor neuron output. First, we found Err2/3 to be broadly expressed by all gamma motor neurons. Second, consistently, gamma motor neuron biophysical properties were disrupted as a whole upon loss of Err2/3. Third, the range of movements affected by Err2/3 removal from motor neurons (from running to ‘skilled’ movement) further suggest that Err2/3 function would be required for both static and dynamic modulation of spindle function. The mechanistic bases underlying the two different outputs of gamma motor neurons to spindles thus remain to be addressed. How about the beta motor neurons? Beta motor neurons connect to both extrafusal and intrafusal muscle fibers but are otherwise morphologically indistinguishable from alpha motor neurons^34-36^. Since beta motor neurons, like the alpha motor neurons, possess relatively large soma sizes and receive monosynaptic Ia afferent input^36^, we were able to rule out that Err2/3 were also operating in beta motor neurons. Since beta motor neurons so far have mostly been studied in the cat, and have been identified solely based on their simultaneous innervation of intrafusal and extrafusal fibers^34-36^, both the prevalence of beta motor neurons and their significance for movement control in rodents awaits further study.

The remarkable contribution of a subset of motor neurons to proprioceptive movement control in mammals has long been recognized^4-6^, but in over 75 years since the description of gamma motor neurons, little has been learned about how these neurons acquire properties that allow them to function differently from the force-generating motor neurons in the first place. In the present study, we identified a long-sought mechanism driving the functional divergence of gamma and alpha motor neurons, by showing that the orphan nuclear receptors Err2/3 implement a ‘feature-specific’ terminal differentiation program promoting the acquisition of gamma motor neuron functional properties (Figure 7E). These properties, in turn, allow the nervous system to engage muscle spindles for regulating muscle proprioception and thus movement accuracy. The action of Err2/3 in gamma motor neuron functional specification could ultimately serve as a blueprint for how such feature-specific terminal differentiation programs impart either functional or structural properties elsewhere in the nervous system.

## Experimental Procedures

### Mouse models

Mouse husbandry and experiments involving mice were approved by and conformed University, state, federal and European Union animal welfare regulations. For experiments using wild-type mice, C57BL/6J and CD1 strains (both purchased from Charles River Laboratories, Inc., Wilmington, USA) were used. “*Chat::*tdTomato” mice were generated by interbreeding mice carrying “*Rosa26*^*floxtdTomato*^*”*^*37*^ and *“Chat*^*Cre*^*”*^*38*^ targeted alleles (both purchased from Jackson Laboratories, Bar Harbor, USA). Egr3^ko/ko^ mice^39^ were a gift from Warren G. Tourtellotte (Northwestern University, currently Cedars Sinai Medical Center). Err2/3^cko^ (*Esrrb*^*flox/flox*^; *Esrrg*^*flox/flox*^; *Chat*^*Cre*^) mice were obtained by interbreeding mice carrying *Chat*^*Cre*^ and floxed *Esrrb*^*loxp/loxp*^ (exon 2 of *Esrrb* gene (chromosome 12) has been flanked by two loxP sites)^40^ alleles with mice carrying floxed *Esrrg*^*loxp/loxp*^ (in which exon 2 of *Essrg* gene (chromosome 1) has been flanked by two *loxP* sites alleles, and were derived from the ES cell repository of the Institut Clinique de la Souris (ICS), Alsace, France). Unless otherwise indicated, “controls” for comparison with Err2/3^cko^ mice were of the genotype *Esrrb*^*flox/flox*^; *Esrrg*^*flox/flox*^ (unrecombined alleles) and Egr3^ko/+^ or C57bl6/J (wild type) mice for comparison with Egr3^ko/ko^ mice.

### Immunodetection

30-60 μm frozen OCT (Sakura Finetek GmbH, Umkirch, Germany) sections were incubated overnight in PBS containing 1% BSA and 0.5% Triton-X 100 as described^41-43^, using the following antibodies: mouse anti-NeuN (1:1500, Cat. # MAB377, Merck KGaA, Darmstadt, Germany), chick anti-GFP (1:2000, Cat. # ab13970, Abcam plc., Cambridge, UK), rabbit anti-dsRed (1:1000, Cat. # 632496, Takara Bio Europe SAS, Saint-Germain-en-Laye, France), mouse anti-Err2 IgG2b (1:4000, Cat. # PP-H6705-00, R&D Systems Inc., Minneapolis, USA), mouse anti-Err3 IgG2a (1:1000, Cat. # PP-H6812-00, R&D Systems Inc., Minneapolis, USA), rabbit anti-VAChT (1:750, Cat. # 139103, Synaptic Systems GmbH, Göttingen, Germany), guinea pig anti-VGLUT1 (1:1000, Cat. # AB5905, Merck KGaA, Darmstadt, Germany), rabbit anti-Isl1/2 (1:2500, Gift from S. L. Pfaff, Salk Institute La Jolla, CA USA), mouse anti-V5 (1:1000, Cat. # 37-7500, Thermo Fisher Scientific Inc., Waltham, USA). Secondary detection of anti-Err2 and anti-Err3 IgG isoforms was accomplished by Alexa Fluor-conjugated anti-mouse IgG2a (1:2000, Cat. # A-21131, Thermo Fisher Scientific Inc., Waltham, USA) or IgG2b (1:2000, Cat. # A-21147, Thermo Fisher Scientific Inc., Waltham, USA) antibodies.

### Microscopy, image analysis and quantification

Fluorescence microscopy was performed using Zeiss LSM 710 and LSM 800 laser scanning microscopes. 6-28 optical sections were obtained at a step-size of 0.8-1.5 μm. Care was applied to avoid oversaturation and distortion of relative expression levels during image acquisition. Raw images were imported into ImageJ and z projected at maximum intensity. For quantification of expression levels raw pixel intensities were quantified in individually outlined motor neuron nuclei “region of interests” (ROIs) using Adobe Photoshop CS5.1. using background levels and the neuron with the highest fluorescent intensity for normalization. For quantification of VGLUT1^+^ varicosities, a step-size of 0.80 μm was used to obtain an average of 20 optical sections. Raw Z-stack Carl Zeiss files (.czi) were imported into *Imaris 8*.*0* (Bitplane AG, Zurich, Switzerland). Neuronal surfaces were rendered to detect vGlut1 varicosities (“spots”) on motor neuron somas and dendrites using the “find spots close to surface” function^44^ and guided by specific parameters^24^.

### Electrophysiology of gamma vs. alpha/beta motor neurons in mice

Mice (P20-22) were intraperitoneally injected with 0.5%-2% (w/v) FluoroGold (FG, Flurochrome LLC., Denver, CO) dissolved in PBS (pH=7.2) at a volume of 0.10 ml/10 g body weight. The animals (1-day post-FG injection) were intraperitoneal injected with 100 mg/kg body weight of Ketamine, 20 mg/kg body weight Xylazine in PBS pH=7.2 at a volume of 0.10 ml/10 g of body weight. After losing their righting reflex, they were placed on a bed of ice until loss of toe pinch response. Immediately after, the animals were decapitated and quickly eviscerated. The torso was placed in chilled Dissecting aCSF (DaCSF) solution (in mM): 191 sucrose, 0.75 K-gluconate, 1.25 KH_2_PO_4_, 26 choline bicarbonate (80% solution), 4 MgSO_4_, 1 CaCl_2_, 20 dextrose, 2 kynurenic acid, 1 (+)-sodium L-ascorbate, 5 ethyl pyruvate, 3 myo-Inositol. The solution was maintained at pH ∼7.3 using carbogen (95% O_2_-5% CO_2_), and osmolarity was adjusted to ∼305-315 mOsm with sucrose. Vertebrectomy was performed to extract the spinal cord. Ventral roots were cut and the meninges were removed from the spinal cord. The thoracolumbar region (T10-L5) of the spinal cord was isolated and embedded in agar block (4% agar in Recording aCSF (RaCSF)) using 20% gelatin in RaCSF). Slices (370 μm) were obtained using Leica VT1200 S (Leica Biosystems, GmbH, Nussloch Germany). The slices were incubated in 35°C RaCSF for 30 minutes and 30 minutes at room temperature before the recordings. Motor neurons (MNs) were recorded in the RaCSF solution (mM): 121 NaCl, 3 KCl, 1.25 NaH_2_PO_4_, 25 NaHCO_3_, 1.1 MgCl_2_, 2.2 CaCl_2_, 15 dextrose, 1 (+)-sodium L-ascorbate, 5 ethyl pyruvate, 3 *myo*-Inositol. The solution was maintained at pH ∼7.4 using carbogen (95% O_2_-5% CO_2_), and osmolarity was adjusted to ∼305-315 mOsm with sucrose. Whole cell patch-clamp recordings were performed from FG-labeled motor neurons in the ventral horn from control and Err2/3^cko^ animals. FG^high^ and FG^low^ MNs were visually identified using Olympus BX51W1 microscope (Olympus Europa SE & Co. KG, Hamburg, Germany) equipped with an FG longpass filter set (350nm bandpass and 425nm longpass filter) (AHF analysentechnik AG, Tübingen Germany). The patch pipette (resistances of 3-6 MΩ) was filled with intracellular solution (mM): 131 K-methanesulfonate (or MeSO_3_H), 6 NaCl, 0.1 CaCl_2_, 1.1 EGTA-KOH, 10 HEPES, 0.3 MgCl_2_, 3 ATP-Mg^2+^ salt, 0.5 GTP-Na^+^ salt, 2.5 L-glutathione reduced, 5 phosphocreatine di(tris) salt. The solution pH was adjusted to 7.25 with KOH and the osmolarity was adjusted to 300 mOsm using sucrose. Data analysis was performed offline using Axograph X Version 1.6. Previously established protocols were applied to obtain membrane properties of rheobase, input resistance, capacitance, *F-I* curve, AHP amplitude, half-width and half-decay times^21, 45-48^. For obtaining the *F-I* curve for discharge properties, spikes were elicited by applying 20 pA, 1000 ms square current pulses to cells. Currents up to 1 nA were injected for all neurons. For the mouse recordings, currents of up to 1 nA were injected into FG^high^ and 3 nA into FG^low^ MNs from control and Err2/3^cko^ spinal cord slices. The firing frequency (Hz) was defined as the inverse of the duration between first two spikes (instantaneous firing frequency), or 0.25-0.75 seconds of the current pulse (steady-state firing frequency), or 1 second current pulse (mean firing frequency). The gain (Hz/nA) was defined as the slope of the regression line of mean firing frequency upon current injection^49,65^.

### Gait analysis

Control (n =9) and Err2/3^cko^ (n=8) mouse locomotion on a treadmill was recorded through automated high-speed motion-capture (DigiGait, Mouse Specifics Inc., Framingham, USA) as described previously^21, 50, 66^. This method generates over 50 different gait variables, which exceed the number of observations (8-9 animals per genotype) and are partially redundant. We therefore used the Partial Least Squares (PLS) regression which is optimized for predictive modeling of multivariate data and to deal with multicollinearity among variables. Orthogonal Signal Correction PLS (OSC-PLS) was used as an extension of PLS to separate continuous variable data into predictive and uncorrelated information for improved diagnostics as well as more easily interpreted visualization. The method seeks to maximize the explained variance between-groups in a single dimension and separates the within-group variance (orthogonal to classification goal) into orthogonal dimensions or components. We modified an existing R script^51,52^, originally designed for chemometrics analysis^53^, to adapt it to our behavior data. The OSC-PLS method was applied to the complete centered-scaled dataset in order to define a model. This highly complex model was then optimized to establish a robust model, the most parsimonious with the highest prediction performance (not shown). A model with an optimal number of 2 components was used for subsequent analysis in both fore and hindlimbs. Our model coefficient of determination Q2, i.e. the model’s fit to the training data, and its Root Mean Square Error of Prediction (RMSEP), i.e. the model’s predictive ability on the testing data were calculated using the Leave-One-Out method^51^. The model was finally validated to ensure that it was performing better than a random model, while not being over-fitted. We conducted an internal cross validation by performing permutations in the original data, from which 2/3 was used to fit a model. This model was then used to predict group memberships on the remaining 1/3 testing set. The process was repeated 100 times and Q2 and RMSEP values were averaged over the repeats. We finally compared our model’s Q2 and RMSEP values to the mean Q2 and mean RMSEP values of the permuted models. The results of the two-sample Student’s t-tests used for the comparisons indicated a probability much inferior to 0.1% of achieving a performance similar to our model by random chance.

### Precision Movement Tasks

Precision movements were successively tested using a custom-built setup with a 100 cm horizontal beam of 20, 25 and 30 mm width, respectively^54^. Age-matched control (n=4) and Err2/3^cko^ (n=4) 10-month-old female mice were trained to move across the beam into a home cage at its end. The animals were trained for 3 days (4 trials/animal, bidirectional on the beam) and tested on the fourth day (4-5 trials/animal, bidirectional on the beam). Furthermore, a custom-built setup to record skilled locomotion on a 100 cm horizontal ladder, with 3 mm rungs spaced at 14 mm (similar to the setup previously described)^9^ was used to test age-matched 8 weeks-old female control (n=5) and Err2/3^cko^ (n=5) mice. Mice were trained to move across the ladder into a home cage placed at its end and were trained for 2 days (3 trials/animal, single direction on the ladder) for two-weeks and tested on the third day of the second week (4-5 trials/animal, single direction on the ladder). The animals were rested in their home cage for 1 minute between trials for both tasks. Animal locomotion was recorded using GoPro HD Hero2 (GoPro Inc., San Mateo, U.S.A) fitted to a custom-built slider track. The videos were acquired at 120 fps at an image size of 848×480 and stored as MP4 files. The videos were processed using GoPro Studio Version (2.5.4) and proDAD Defishr Version 1.0 (proDAD GmbH, Immendingen, Germany). The figure videos were slowed to 25-40% of original speed and reduced to 60 fps using GoPro Studio Version (2.5.4). The fish-eye view was removed from videos using proDAD Defishr Version 1.0 (proDAD GmbH, Immendingen, Germany) with Mobius A Wide presets and Zoom adjusted to 110.0 or 180.0. A “miss” was scored when the mouse paw failed to locate the rung or the beam leading to the animal to slip or to halt until the paw regained its footing. An average of ∼40 steps per trial were analyzed for each mouse.

### Muscle spindle afferent recordings

*Ex vivo* extensor digitorum longus (EDL) muscle-nerve preparations were used to study the response of muscle sensory neurons to stretch and was essentially performed as described^31^. Briefly, the dissection of muscle (with the nerve attached) was performed in low calcium and high magnesium solution^31,55^. Then, the muscle was placed in recording solution 22-24 °C and equilibrated with carbogen and was then hooked to a dual force and length controller -transducer (300C-LR, Aurora Scientific Inc., Aurora, Canada) with the help of 5-0 sutures tied to its tendons. Following the determination of resting length (Lo) as described in previous studies, a suction electrode (tip diameter 50-80 μm) connected to an extracellular amplifier (EXT-02F, npi Electronics GmbH, Tamm Germany) was used to sample muscle spindle afferent activity. Data acquisition was performed with LabChart 8 connected to PowerLab 8/35 (ADInstruments Ltd., Oxford, UK). Afferent activity, when obtained was checked for the presence of a characteristic pause following a series of 30 twitch contractions at 1Hz and if the pause was present, the unit was identified as a muscle spindle afferent. Then a series of 9 ramp and hold stretches were delivered to the muscle (2.5% Lo, 5% Lo and 7.5% Lo at 40% Lo/sec, protocols were kindly provided by K.A. Wilkinson lab). The data was recorded and analyzed offline with a custom written MATLAB code. Spikes were detected using KMEANS (2). For each afferent, resting discharge (RD), dynamic peak discharge (DP), dynamic index (DI) and static response (SR) were calculated. A total of 10 control and 10 Err2/3^cko^ animals were used for recordings, from which 8 control and 9 Err2/3^cko^ spindle afferents were analyzed.

### Molecular cloning

Mouse *Esrrb* (NM_011934.4) (Err2) and Mouse *Esrrg* (NM_011935.3) (Err3) open reading frames were isolated using PrimeScript 1st cDNA synthesis Kit Takara Bio Europe SAS, Saint-Germain-en-Laye, France) following manufacturer’s directions from E18.5 mouse spinal cord total RNA and cloned after PCR amplification with the following oligonucleotide primers:

*Esrrb* Forward 5’-CATGCCATGGATGTCGTCCGAAGACAGGCACC-3’,

*Esrrb* Reverse 5’-CATGCCATGGCACCTTGGCCTCCAGCATCTCCAGG-3’,

*Esrrg* Forward 5’-CATGCCATGGATGGATTCGGTAGAACTTTGC-3’,

*Esrrg* Reverse 5’-CATGCCATGGGACCTTGGCCTCCAGCATTTCC-3’.

The chick *Kcna10* (NP_989793) open reading frame was isolated from E5.5 chick embryo total RNA as above using the following primers:

*Kcna10* Forward 5’-ATGATGGACGTGTCCAGTTGG-3’

*Kcna10* Reverse 5’-TTTTTTGGCCTTGTCTCGAGG-3’.

Thus, synthesized cDNAs were subcloned into an expression vector between *CMV* promoter in frame with an Aphthovirus 2A peptide-GFP fusion sequence for co-translational cleavage^56^. This entire cassette was flanked by *Tol2* sites facilitating genomic integration upon co-transfection with Tol2 transposase as described^21^. Err2VP16 was generated by fusing the open reading frame for the herpes simplex virus-1 (HSV-1) VP16 (amplified from *pActPL-VP16AD* plasmid, Addgene plasmid #15305, Watertown, USA) activation domain to Err2. Err2EnR was generated by fusing the open reading frame of the transcriptional repressor domain of *Drosophila* Engrailed (amplified from *CAG-EnR* plasmid, Addgene plasmid #19715, Addgene, Watertown, USA) to Err2.

### Chick motor neuron electrophysiology

Chick embryos electroporated at E2.7-E3.0 (HH stages 14-18) with appropriate DNA constructs were harvested at E12-15 (HH stages 38-41) and processed as described previously^21^. Briefly, chick embryos were placed on ice for 5 minutes, extracted from the egg, decapitated and dissected in a petri dish containing cold chick aCSF (CaCSF) solution (mM): 139 NaCl, 3 KCl, 1 MgCl_2_, 17 NaHCO_3_, 12.2 dextrose, 3 CaCl_2_. The solution pH was adjusted to 7.25 with KOH and the osmolarity was adjusted to ∼315 mOsm using sucrose. The thoracolumbar region of the spinal cord (with the vertebral column intact) was isolated and embedded in an agarose block (4% agarose in CaCSF) using 20% gelatin in CaCSF. Slices (370 μm) were obtained using a Leica VT1200 S vibrating blade microtome (Leica Biosystems GmbH, Nussloch, Germany) and incubated in CaCSF for 30 minutes at room temperature (22°C). Motor neurons were visualized by GFP expression using 4x air objective (Olympus UPlan FL N) and 40x water-immersion objective (Olympus UPlan FI N) equipped on an Olympus BX51W1 microscope. The patch pipette (resistances of 3-6 MΩ) was filled with intracellular solution (mM): 130 MeSO_3_H, 10 KCl, 2 MgCl_2_, 0.4 EGTA, 10 HEPES, 2 ATP-Mg^2+^ salt, 0.4 GTP-Na^+^ salt, 0.1 CaCl2. The solution pH was adjusted to 7.3 with KOH and the osmolarity was adjusted to ∼315 mOsm using sucrose. The intracellular solution contained 25 μM Alexa fluor 568 dye (Thermo Fisher Scientific, Inc., Waltham, USA) to label recorded motor neurons. Current-clamp recording signals were amplified and filtered using MultiClamp 700B patch-clamp amplifier (Molecular Devices LLC., San Jose, USA). The signal acquisition was performed at 20 kHz using Digidata 1322A digitizer (Molecular Devices LLC., San Jose, USA) and pCLAMP 10.4 software (Molecular Devices LLC., San Jose, USA).

### RNA sequencing and transcriptome analysis

E12.5 (HH St. 38-39) chick lumbar spinal motor columns transfected with either *CMV::eGFP*, or *CMV:: Err2VP16*.*2A*.*eGFP* or *Dlk1*.*IRES*.*eGFP* plasmids were identified by GFP fluorescence, dissected and collected. RNA isolation and RNA sequencing were carried out as described previously^57^. Briefly, RNA was isolated using Tri-Reagent (Sigma-Aldrich Chemie GmbH, Taufkirchen, Germany) and Phenol-Chloroform extraction according to the manufacturer’s protocol. RNA quality was assessed using Nanodrop 2000 (Thermo Fisher Scientific, GmbH) and RNA integrity number (RIN) was evaluated by using the Agilent 2100 Bioanalyzer (Agilent Technologies, Inc., USA). RNA was reverse transcribed to cDNA using Transcriptor High Fidelity cDNA synthesis kit (Roche Diagnostics Deutschland GmbH, Mannheim, Germany) and RNA-Seq libraries were obtained using TruSeq RNA Sample Preparation v2 kit (Illumina, Inc., San Diego, USA). To analyze the library quality, Agilent 2100 Bioanalyzer (Agilent Technologies, Inc., Santa Clara, USA) was used and the concentration was measured by a Qubit^R^ dsDNA HS Assay kit (Thermo Fisher Scientific Inc., Waltham, USA). The concentration was adjusted to 2 nM prior to sequencing (50 bp) on a HiSeq 2000 sequencer (Illumina, Inc., San Diego, USA) using TruSeq SR Cluster kit v3-cBot-HS (Illumina, Inc., San Diego, USA) and TruSeq SBS kit v3-HS (Illumina, Inc., San Diego, USA) based on manufacturer’s instructions. RNA-sequencing quality was evaluated by utilizing raw reads using the FastQC quality control tool version 0.10.1^58^. Bowtie2 v2.0.2 using RSEM version 1.2.29 with default parameters was utilized to align sequence reads (single-end 50 bp) to chicken reference genome (Galgal5)^59, 60^. Prior to indexing, GFP, Err2, Dlk-1, VP16 and IRES sequences and annotations were added to the reference genome (FASTA file) and annotations (GTF file). Ensembl annotations (version 86.5) with rsem-prepare-reference from RSEM software was used to index chicken reference genome^61^. Furthermore, sequence alignment of sequence reads and gene quantity was obtained through the use of rsem-calculate-expression. Rsem-calculate-expression resulted in sequence read count and TPM value (transcripts per million) for individual genes. DESeq2 package was used to carry out differential expression analysis^62^. Finally, genes with less than 5 reads (baseMean) were filtered, while genes with an adjusted p-value < 0.05 were classified as differentially expressed. Gene ontologies and categorization was performed using the DAVID Gene Functional Classification Tool^67^.

### Enhancer identification and promoter assays

Evolutionary conserved non-coding genomic regions (ECRs) around the *Kcna10* genomic locus were identified using the ECR Browser^63^ and screened for potential Err2/Err3 transcription factor binding sites using the JASPAR CORE database^64^. A 240 bp ECR 3.5 kb upstream of the *Kcna10* transcription start site with three putative Err2/Err3 binding sites was amplified from mouse genomic DNA using the following primers: Forward 5’-TCTCACAGCCCTGCTCATC-3’ and Reverse 5’-CTTGCCTGAGAACCTGATCTCC-3’ and subcloned into a reporter vector containing a minimal promoter followed by tdTomato coding sequence, which together were flanked by *Tol2* sites to facilitate stable genomic integration. To test promoter activity and potential regulation by Err2/3, Lohmann LSL fertilized chick eggs were incubated until E2.7-E3.0 (HH stages 14-18) and chick embryo neural tubes were electroporated *in ovo* using the ECM 830 electroporation system (BTX/Harvard Apparatus Inc., Holliston, USA) as described^21^. *Kcna10::tdTomato* reporter plasmids together with either *CMV::2A*.*eGFP* or *CMV::VP16:Err2*.*2A*.*eGFP* at a molar ratio of 1:1 were transfected.

## Data Availability

All data are available in the manuscript or the supplementary materials. Raw data are available upon reasonable request to the corresponding author.

## Acknowledgments

We thank Louisa Dorr, Simeon Helgers, Beate Veith, Alexandra Klusowski, Brenda Ross for technical assistance and Rob Brownstone (UC London) for help setting up P20-22 motor neuron recordings, Warren G. Tourtellotte (Northwestern University, currently Cedars Sinai Medical Center) for the *Egr3*^*ko*^ mice and Stephan Kröger (LMU Munich) for help setting up muscle spindle recordings. This research received funding from the European Research Council under the European Union’s Seventh Framework Programme (FP/2007-2013)/ERC Grant Agreement 311710-MU TUNING, the Göttingen Excellence Cluster for Nanoscale Microscopy and Molecular Physiology of the Brain, Deutsche Forschungsgemeinschaft (DFG) Research Center 103, Section B1 and the BMBF. Ashish Rajput, Vikas Bansal, and Stefan Bonn were supported by BMBF IDSN, ERA-Net E-Rare MAXOMOD, and CRC 1286/Z2. Dario Farina received funding from the European Research Council through the Synergy Grant NaturalBionicS (contract #810346). Francesco Negro has received funding from the European Union’s Horizon 2020 research and innovation programme under the Marie Skłodowska-Curie grant agreement No 702491 (NeuralCon).

## Author Contributions

M.N.K., P.C., F.N., A.R., P.F., T-I.L. Y.B. and W.P.M. conducted the experiments, M.N.K., P.C., F.N., V.B., C.L., D.M., T.A., S.B. and D.F. analyzed the data, M.N.K. and T.M. designed the experiments and wrote the paper.

