## Supplementary information for "Orphan nuclear receptors Err2 and 3 promote a feature-specific terminal differentiation program underlying gamma motor neuron function and proprioceptive movement control"

**This PDF file includes:**

Supplementary Figs. S1 to S9
Supplementary Tables S1 to S3

**Other Supplementary Materials for this manuscript include the following:**

Supplementary Movies S1 to S4

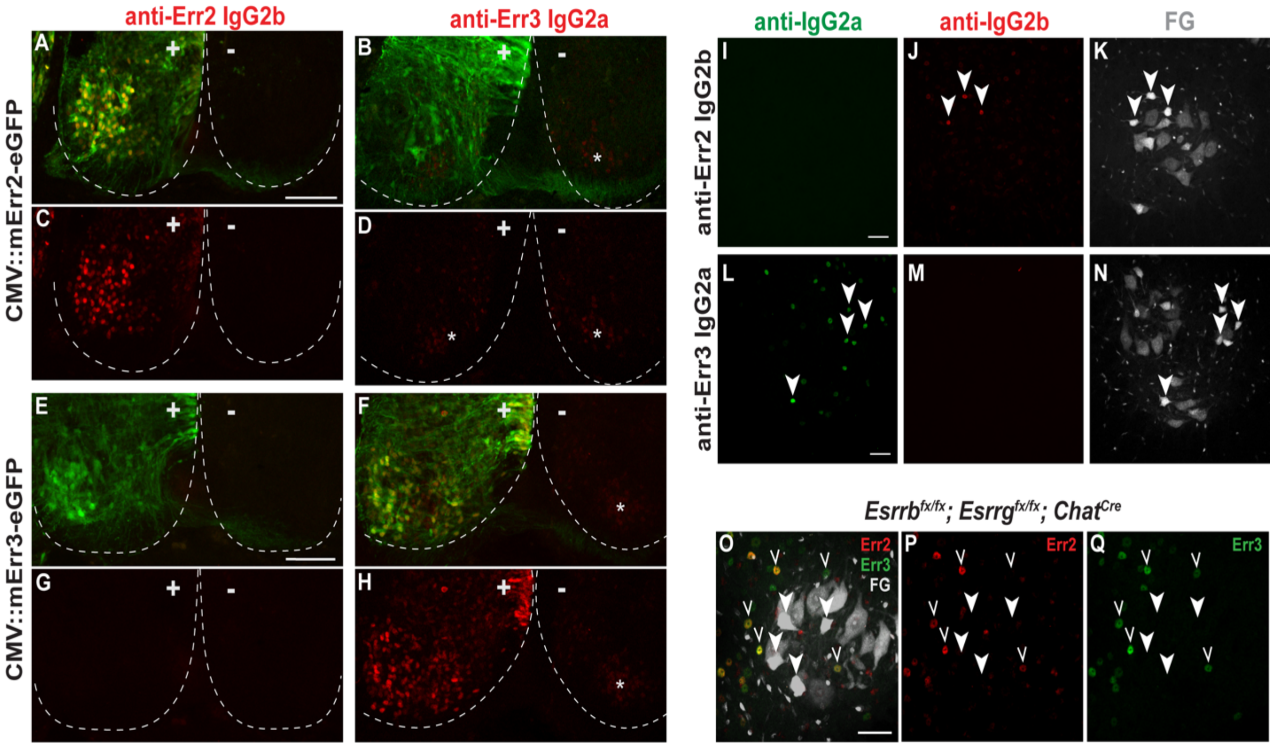


**Figure S1.** Selective immunodetection of either Err2 or Err3.

**(A-D)** Transversal sections of E6 chick spinal cords unilaterally transfected by murine Err2 (mErr2) and eGFP. **(A, C)** Immunofluorescence with anti-Err2 IgG2b detects transfected mErr2 but not endogenous chick Err2 (cErr2) (scale bar: 100 µm). **(B, D)** Anti-Err3 IgG2a detects at low-levels of endogenous cErr3 (asterisks) but does not cross-react with transfected mErr2. **(E-H)** Transversal sections of E6 chick spinal cords unilaterally transfected by murine Err3 (mErr3) and eGFP. **(E, G)** Anti-Err2 IgG2b does not cross-react with mErr3. **(F, H)** Anti-Err3 IgG2a detects transfected mErr3 and also detects at low-levels of endogenous cErr3 (asterisk in contralateral spinal cord) but does not cross-react with transfected mErr2. **(I-N)** Transversal sections of P21 mouse spinal cord ventral horn. **(I)** No cross-reactivity of Alexa488 anti-IgG2a with anti-Err2 IgG2b (scale bar: 50 µm). **(J)** Detection of anti-Err2 IgG2b with Alex555 anti-IgG2b in a subset of motor neuron nuclei (arrowheads, note: lower levels detected in other motor neuron and interneuron nuclei). **(K)** FluoroGold (FG)-traced motor neurons. Note: characteristic high-levels of FG incorporation in small soma size motor neurons (arrowheads). **(L)** Detection of anti-Err3 IgG2a with Alexa fluor 488 anti-IgG2a in a subset of motor neuron nuclei (arrowheads, note: lower levels detected in other motor neuron and moderate-to-high levels in interneuron nuclei) (scale bar: 50 µm). **(M)** No cross-reactivity of Alexa fluor 555 anti-IgG2b with anti-Err3 IgG2a. **(N)** FG-traced motor neurons. **(O-Q)** Transversal sections of P21 Err2/3$^{cko}$ (*Esrrb$^{flox/flox}$*; *Esrrg$^{flox/flox}$*; *Chat$^{Cre}$*) mouse spinal cord ventral horn. (**O**) FG-traced motor neurons overlaid with Err2 and Err3 immunoreactivity. Closed arrowheads: small soma size FG$^{high}$ motor neurons (scale bar: 50 µm). **(P, Q)** Absence of Err2 **(P)** or

Err3 **(Q)** immunoreactivity in Err2/3$^{cko}$ motor neurons (closed arrowheads). Open arrowheads: Err2 and Err3 expression is retained in interneurons in Err2/3$^{cko}$.

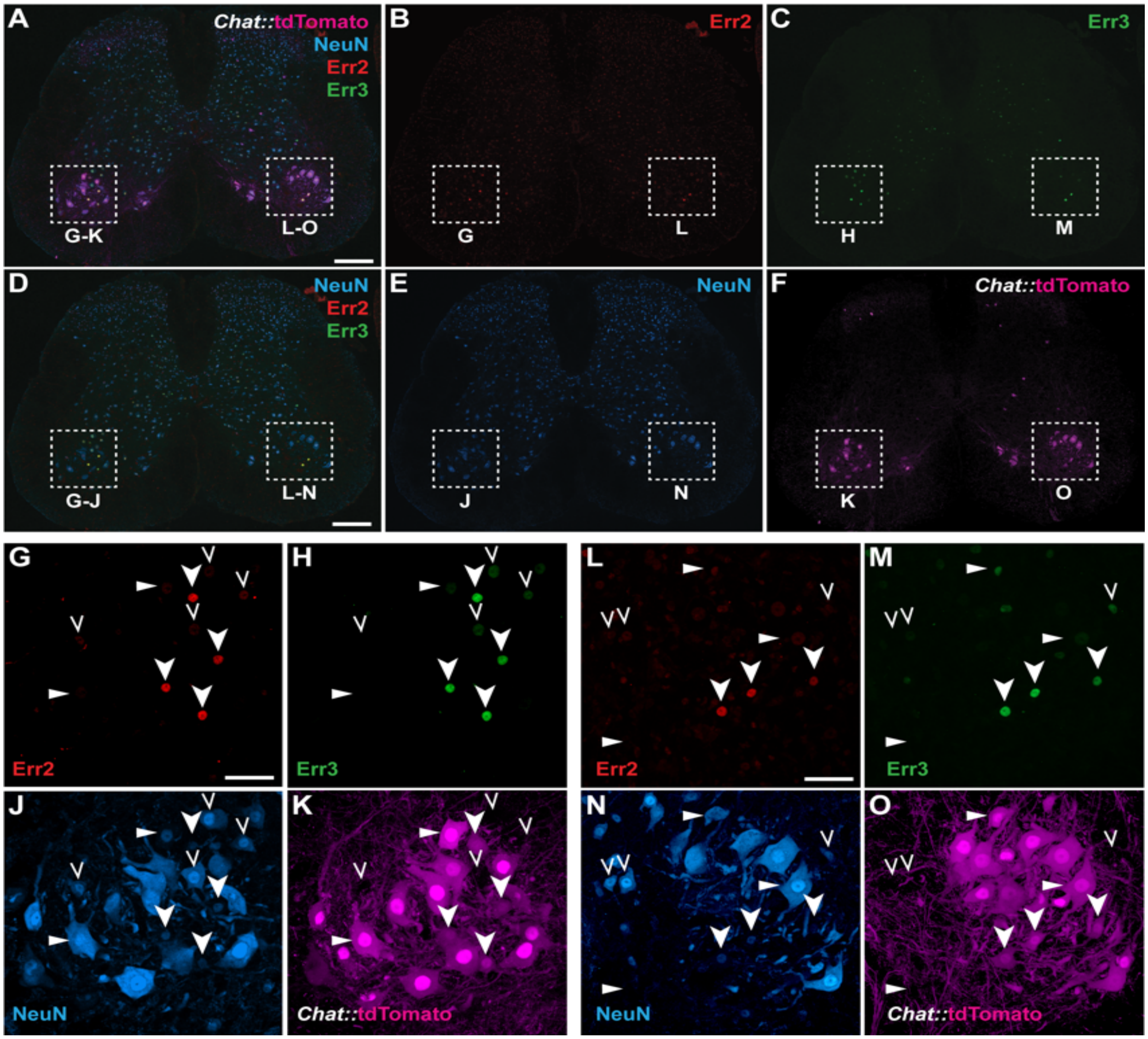


**Figure S2**. High-levels of Err2 and Err3 are co-expressed by gamma motor neurons.

**(A-F)** Overview of transversal section of P21 *Chat::*tdTomato (*Chat*$^{Cre}$; *Rosa26*$^{fxtdTomato}$) lumbar mouse spinal cord. **(A)** Expression of tdTomato, NeuN, Err2 and Err3. tdTomato (magenta) mostly labels motor neurons in lamina IX but also subsets of cholinergic interneurons in laminae III and X (see Figure S4A) (scale bar: 200 μm). High-levels of Err2 and Err3 co-expression in tdTomato$^{+}$ NeuN$^{\text{low or negligible}}$ motor neurons (yellow-orange nuclei in boxed areas). Boxed areas around left and right lateral motor columns correspond to the higher magnification given in G-K and L-O, respectively. **(B)** Highest levels of Err2 in motor neurons (boxed area), relatively lower but detectable levels in other motor neuron subtypes and subsets of interneurons throughout the spinal cord. **(C)** High-levels of Err3 in motor neurons (boxed area), lower but significant levels in other motor neuron subtypes and occasional high-levels in interneurons in the intermediate spinal laminae. **(D)** High-levels of Err2 and Err3 co-expression in NeuN$^{\text{low or negligible}}$ motor neurons (yellow-orange nuclei in boxed areas) (scale bar: 200 μm). **(E, F)** Separate channels depicting expression of NeuN only **(E)** or tdTomato only **(F)**. **(G-K)** Higher magnification of left ventral horn (boxed area) in **(A-F)**. Closed arrowheads: Highest levels of Err2 **(G)** and Err3 **(H)** in consistently NeuN$^{\text{low or negligible}}$ **(J)** and tdTomato$^{+}$ **(K)** small motor neurons (scale bar: 50 µm). Note that small NeuN$^{\text{low or negligible}}$ motor neurons consistently exhibit lower tdTomato levels compared to large motor neurons **(K, O)**. Open arrowheads: low but detectable Err2 **(G)** and Err3 **(H)** levels in tdTomato$^{-}$ interneurons **(K)**. Triangles: low but detectable Err2 **(G)** and Err3 **(H)** levels in large tdTomato$^{+}$ motor neurons **(K)** with intermediate to high NeuN levels **(J)**. Note that NeuN levels vary considerably in large motor neuron subtypes (see Figures 2B and 2F). **(L-O)** Higher magnification of right ventral horn (boxed area) in **(A-F)**. Closed arrowheads: Highest levels of Err2 **(L)** and Err3 **(M)** in consistently NeuN$^{\text{low or negligible}}$ **(N)** and tdTomato$^{+}$ **(O)** small motor neurons (scale bar: 50 µm). Open arrowheads: low but detectable Err2 **(L)** and Err3 **(M)** levels in tdTomato$^{-}$ interneurons **(O)**. Triangles: low but detectable Err2 **(L)** and Er3 **(M)** levels in large tdTomato$^{+}$ motor neurons **(O)** with high NeuN levels **(N)**.

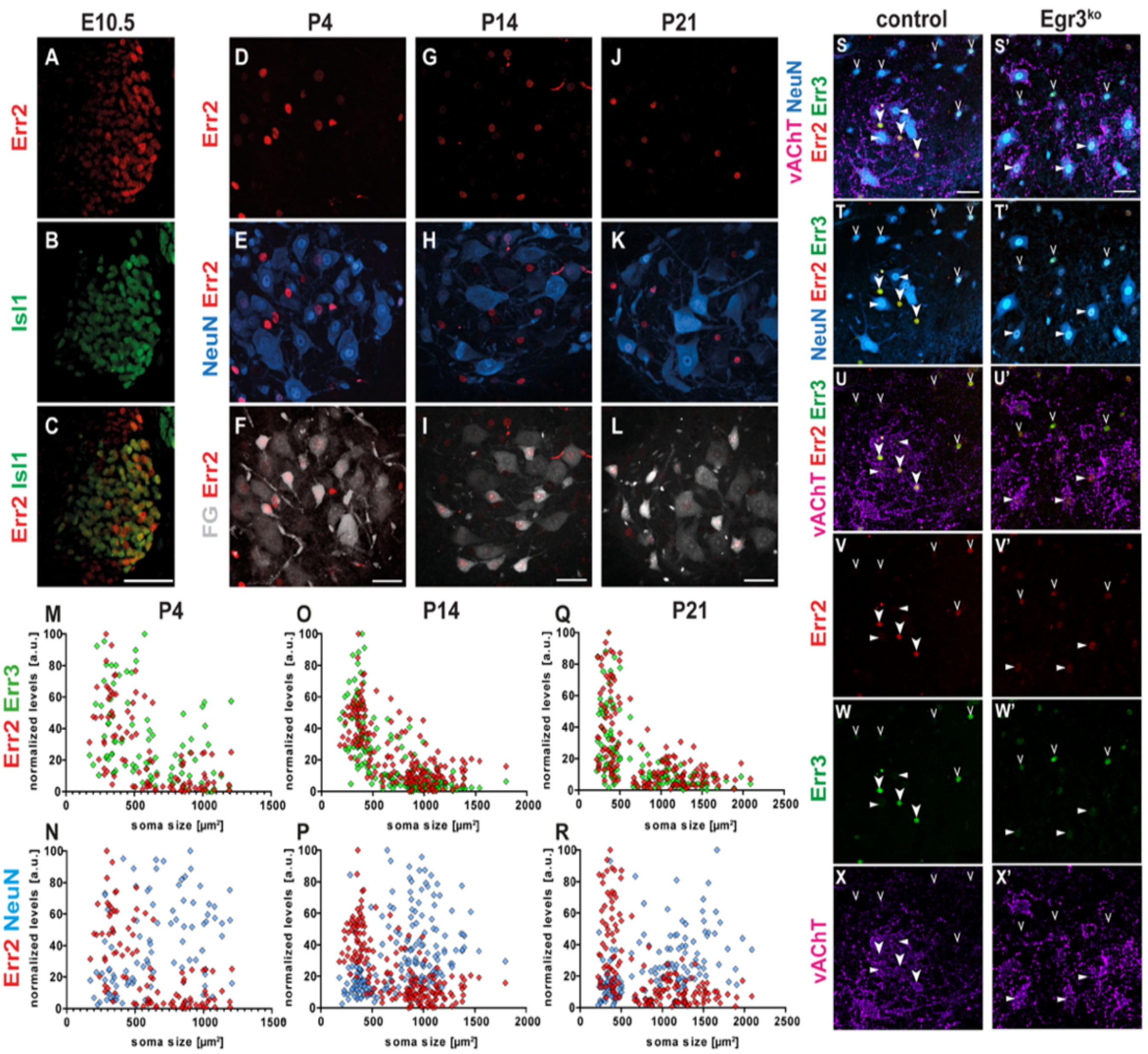


**Figure S3**. Progressive confinement of high Err2 levels to gamma motor neurons during postnatal development depends on muscle spindle-derived signals.

**(A-C)** Expression of Err2 in most Isl1$^{+}$ motor neurons in 10.5 day old mouse embryo thoracic neural tube (scale bar: 50 µm). **(D-F)** At P4 higher levels of Err2 **(D)** have begun to be mostly confined to small NeuN$^{\text{low or negligible}}$ **(E)** motor neurons, while many NeuN$^{\text{high}}$ motor neurons still retain relatively lower yet substantial Err2 levels (compare **D** and **E**). **(F)** FluoroGold (FG) tracing to visualize motor neurons. **(G-H)** At P14 high Err2 levels **(G)** are further confined to small NeuN$^{\text{low or negligible}}$ motor neurons **(H)** with high-levels of FG incorporation **(I)**. **(J-K)** At P21 high Err2 levels **(J)** are further confined to small NeuN$^{\text{low or negligible}}$ motor neurons **(K)** with high-levels of FG retention **(L)**, while Err2 levels in NeuN$^{\text{high}}$ motor neurons have dropped further compared to earlier postnatal stages (compare **D**, **G** and **J**). **(M-Q)** Scatter plots of Err2 (red data points) and Err3 (green) levels over soma sizes show increasing confinement of high-levels of Err2 and Err3 to a population of small motor neurons during postnatal development ages of P4 (n=100), P14 (n=189) and P21 (n=159). **(M-R)** Scatter plots of Err2 (red data points) and NeuN (blue) levels over soma sizes show increasing segregation of motor neurons into $Err2^{high}$, $NeuN^{low\ or\ negligible}$ and $Err2^{low}$, $NeuN^{high}$ populations during postnatal development, while also revealing considerably cell-to-cell variability in relative expression levels of Err2, Err3 and NeuN. **(SS'-XX')** Motor neurons fail to retain high Err2 and Err3 levels in mice with defective spindle development. **(S-X)** Control (transversal section of lumbar spinal cord of adult mouse): expression of high Err2 **(V)** and Err3 **(W)** levels by $vAChT^{+}$ **(X)** $NeuN^{low\ or\ negligible}$ **(T)** motor neurons (closed arrowheads). Open arrowheads: expression of Err2 and Err3 by subsets of $vAChT^{-}$ interneurons. Triangle: low Err2 and Err3 levels in some large motor neurons. Note: $vAChT^{+}$ cytosol identifies motor neurons in lamina IX, while $vAChT^{+}$ varicosities indicate cholinergic synapses or axons stemming from non-motor neurons. **(S'-X')** $Egr3^{ko}$ ($Egr3^{-/-}$) transversal section of lumbar spinal cord of adult mouse: absence of high Err2 **(V')** and Err3 **(W')** levels in $vAChT^{+}$ **(X')** $NeuN^{low\ or\ negligible}$ **(T')** motor neurons. Open arrowheads: retention of Err2 and Err3 expression by $vAChT^{-}$ interneurons (compare **V**, **W** and **X**). Triangles: retention of low Err2 and Err3 levels by large motor neurons (compare **V**, **W** and **X**). n= # of neurons.

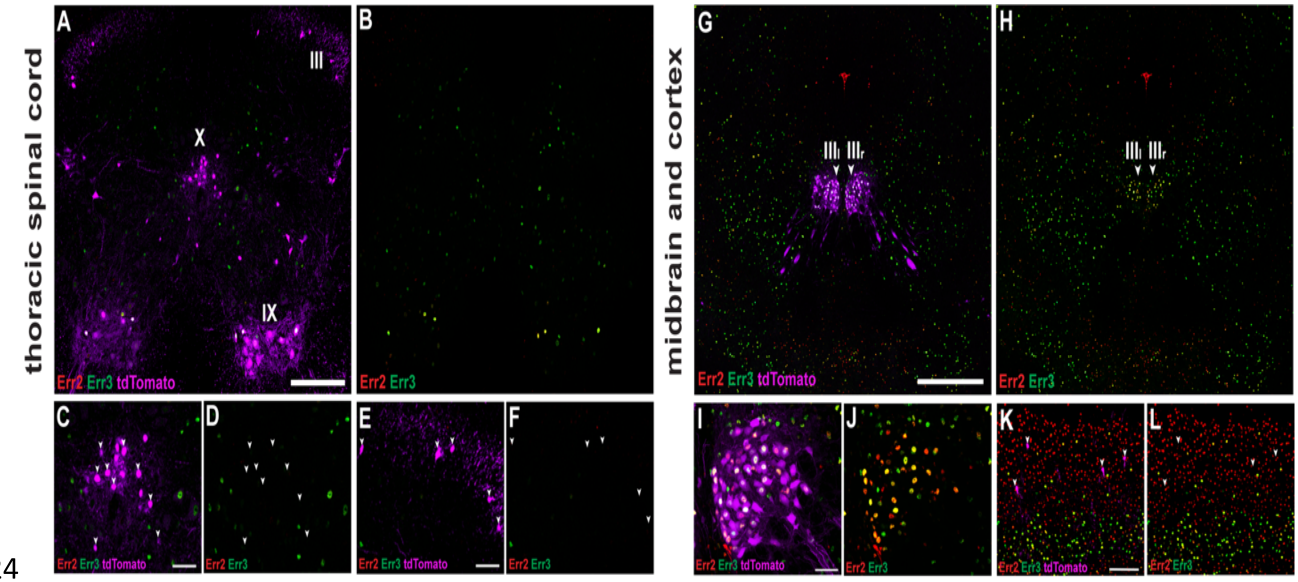


**Figure S4**. Expression of Err2 and Err3 in motor neurons and interneurons.

**(A, B)** Overview of transversal section of P21 *Chat::*tdTomato (*Chat*$^{Cre}$; *Rosa26*$^{fxtdTomato}$) thoracic mouse spinal cord. **(A)** Expression of tdTomato indicating *Chat*$^{Cre}$ activity in motor neurons (lamina IX), interneurons (presumably p0c) in lamina X and lamina III interneurons, as well as few scattered neurons in the intermediate laminae (scale bar: 200 µm). **(B)** Err2 and Err3 expression in tdTomato$^{+}$ motor neurons but not in other tdTomato$^{+}$ neurons outside lamina IX. **(C, D)** No detectable Err2 and Err3 expression in tdTomato$^{+}$ lamina X interneurons (arrowheads) (scale bar: 50 µm). **(E, F)** No detectable Err2 and Err3 expression in tdTomato$^{+}$ lamina III interneurons (arrowheads) (scale bar: 200 µm). **(G, H)** Expression of Err2 and Err3 by oculomotor neurons (III) (arrowheads) and largely non-overlapping expression of either Err2 or Err3 by subsets of non-cholinergic neurons throughout the midbrain (scale bar: 500 µm). **(I, J)** Expression of Err2 and Err3 by most but not all cholinergic neurons of the oculomotor nucleus (scale bar: 50 µm). **(K, L)** Expression of Err2 and Err3 in a subset of non-cholinergic (tdTomato$^{-}$) cortical interneurons (scale bar: 100 µm).

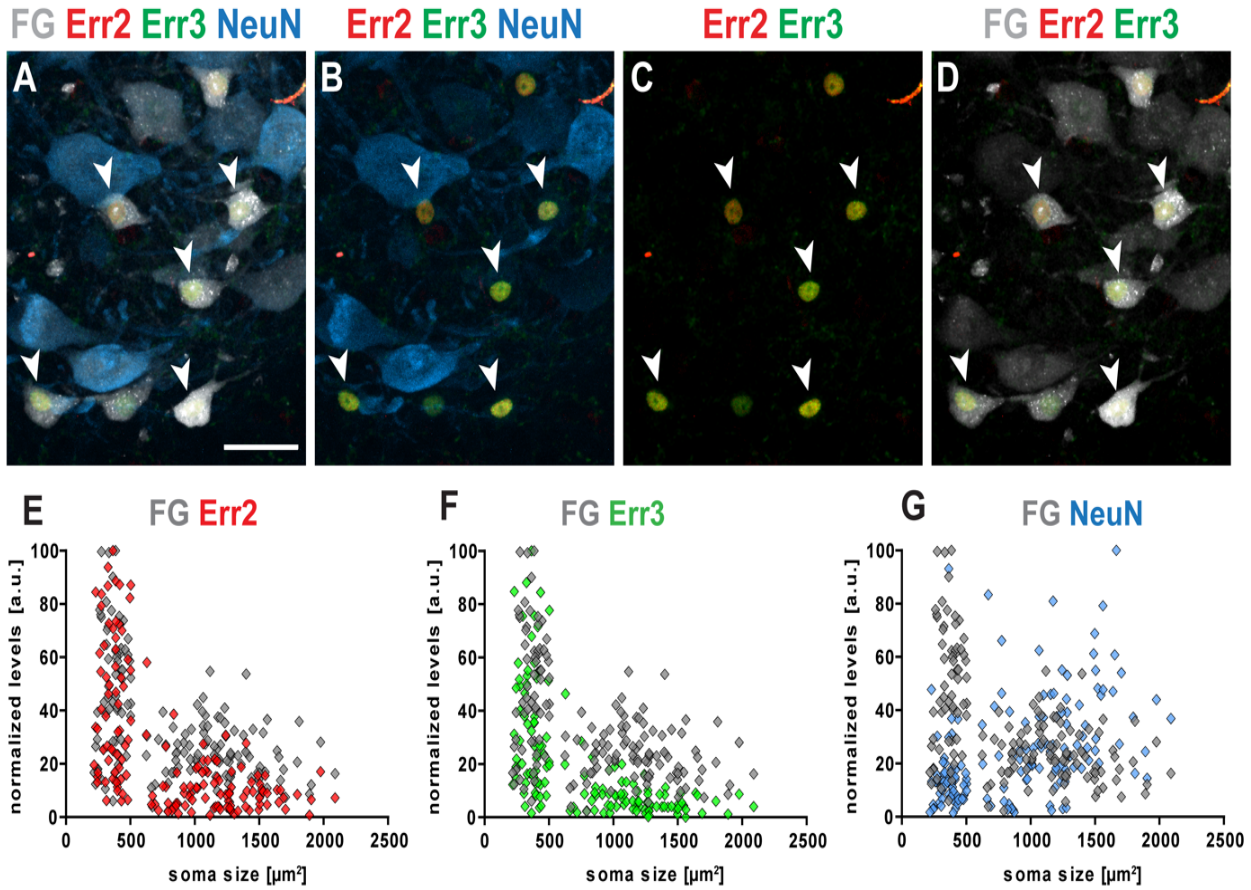


**Figure S5**. Gamma and alpha motor neurons distinguished by FluoroGold (FG) incorporation and soma size at two weeks postnatal.

**(A-D)** P14 spinal cord ventral horn (note: same specimen as in Figure S3G-S3I): gamma and alpha motor neurons can be distinguished by different levels of FluoroGold (FG) incorporation and soma sizes (scale bar: 50 µm). **(A)** Arrowheads: examples of small motor neurons with high-levels of FG incorporation expressing low or negligible levels of NeuN **(B)** and high-levels of Err2 and Err3 **(C)**. **(D)** Different levels of FG in small and large NeuN$^{high}$ motor neurons (compare with **B**). **(E-G)** Scatter plots of motor neuron fluorescence levels over soma sizes at P21 (n=159) (note: data from same experiment depicted in Figure S3Q and S3R): stringent identification of gamma motor neurons based on a combination of soma size and FG levels. **(E, F)** Two distinct populations of motor neurons visualized based on FG levels and soma size: gamma motor neurons with small somas and relatively high FG levels that express high Err2 **(E)** and Err3 **(F)** levels, but low or negligible levels of NeuN **(G)**. Larger motor neurons with lower but substantial FG levels express low or negligible levels of Err2 **(E)** and Err3 **(F)**, but mostly higher levels of NeuN **(G)**.

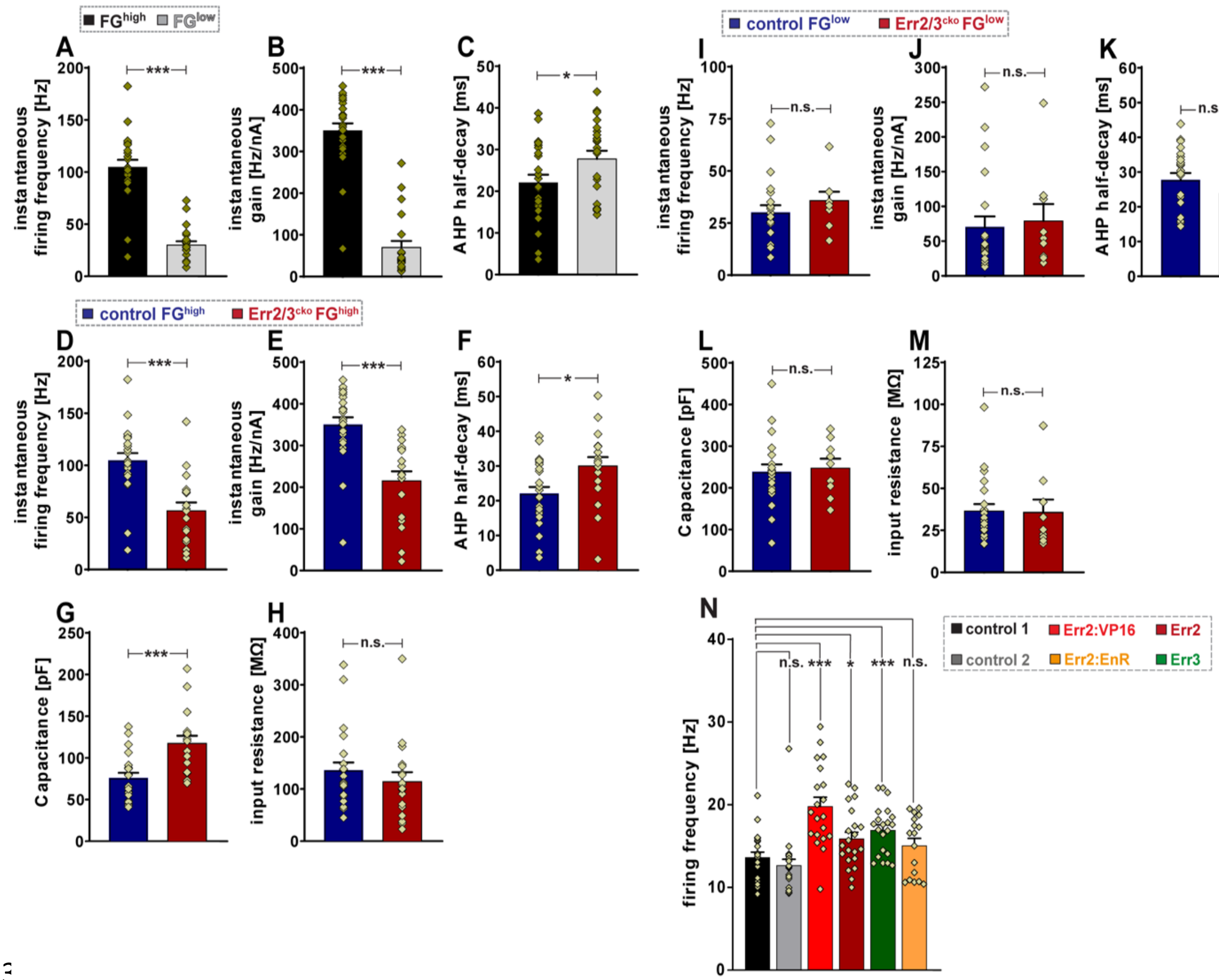


**Figure S6**. Electrophysiological properties of gamma and alpha motor neurons in control and Err2/3$^{cko}$ mice.

**(A-C)** Control FG$^{high}$ (gamma) motor neurons (black bars) exhibit significantly higher instantaneous firing frequency **(A)**, instantaneous gain **(B)** and lower AHP-half decay time **(C)** when compared to FG$^{low}$ (alpha) motor neurons (gray bars). **(D-H)** Err2/3$^{cko}$ FG$^{high}$ (gamma) motor neurons (red bars) exhibit significantly lower instantaneous firing frequency **(D)**, lower instantaneous gain **(E)**, higher AHP-half decay time **(F)**, higher capacitance **(G)**, while no significant difference in input resistance **(H)** when compared to control FG$^{low}$ motor neurons (blue bars), respectively (see Supplementary Table S1 for details). **(I-M)** No significant differences between Err2/3$^{cko}$ FG$^{low}$ (alpha) motor neurons (red bars) and control FG$^{low}$ (alpha) motor neurons (red bars) when comparing instantaneous firing frequency **(I)**, instantaneous gain **(J)**, AHP-half decay time **(K)**, capacitance **(L)** and input resistance **(M),** respectively (see Supplementary Table S1 for details). **(N)** Compared to control 1 (13.63 ± 0.63), significant increase in firing frequency was observed upon over-expression of Err2:VP16 (19.82 ± 1.10), Err2 (15.91 ± 0.78), Err3 (16.93 ± 1.12) but not upon Err2:EnR (15.08

± 0.87) forced expression. Statistically significant differences are indicated as: *p<0.05, **p<0.01, ***p<0.001, n.s.= not significant, Student's t-test).

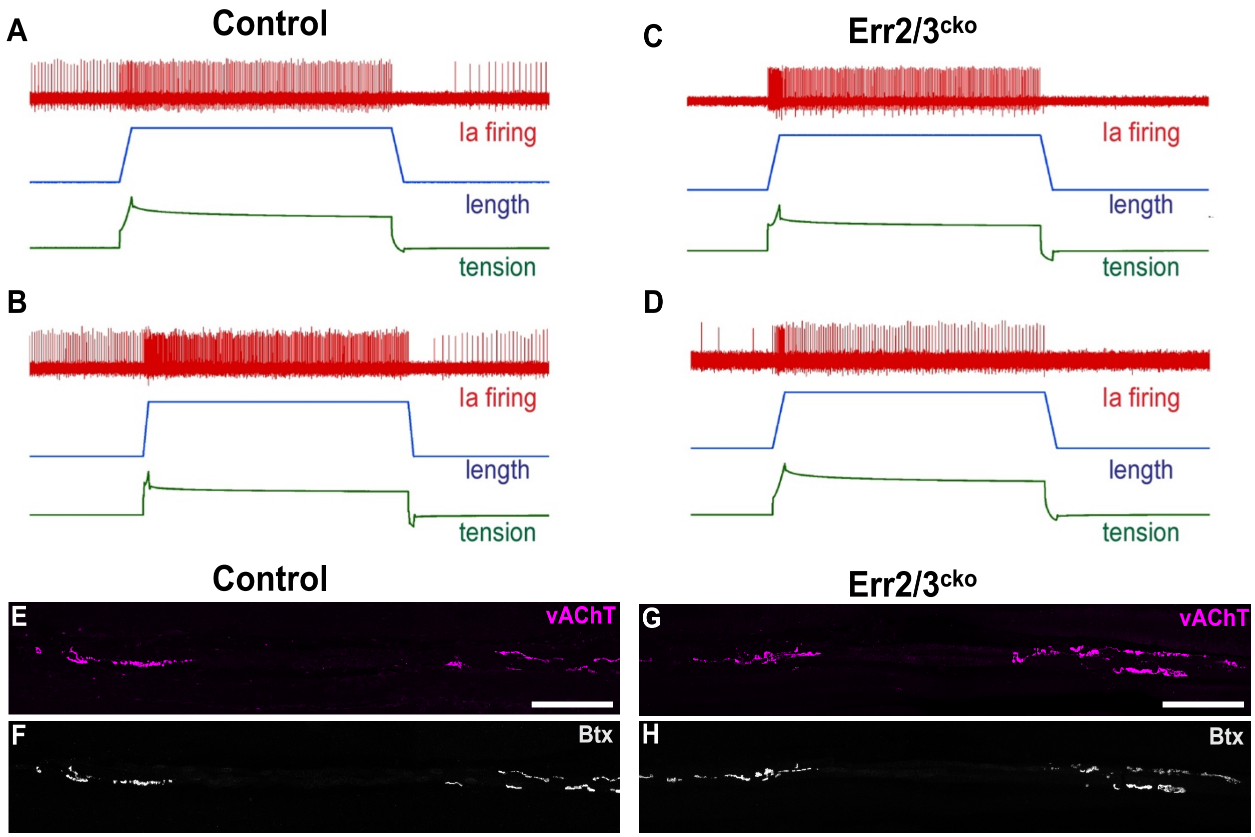


**Figure S7**. **(A-D)** Examples of Ia afferent recordings from extensor digitorum longus (EDL) nerve-muscle preparations. Red traces: Ia afferent firing, blue traces: relative muscle length, green traces: relative muscle tension upon application of muscle stretch with a force transducer.

**(A, B)** In the control preparation, Ia afferents exhibit relatively low tonic firing rates at resting length, which rapidly increases upon application of muscle stretch with two characteristic bursts during stretch onset and offset. **(C, D)** In Err2/3$^{cko}$ mice Ia afferents frequently fall silent during resting length, while exhibiting a responsiveness towards stretch similar to control Ia afferents (compare with **F, H**). **(E-H)** P70 mouse extensor digitorum longus (EDL) muscle spindles of control **(E, F)** and Err2/3$^{cko}$ **(G, H)** mice. **(E-H)** Muscle spindle motor innervation (vAChT, magenta) and their postsynaptic sites (Btx, alpha bungarotoxin, grey) (scale bar: 100 µm). **(E)** Motor axon presynaptic termini within the peripheral contractile segments of the intrafusal muscle fibers. **(F)** Motor endplates associated with the motor axon presynaptic termini within the intrafusal muscle fiber contractile segments. **(G)** Motor axon presynaptic termini within the peripheral contractile segments of the intrafusal muscle fibers (compare with **E**). **(H)** Motor endplates associated with the motor axon presynaptic termini within the intrafusal muscle fiber contractile segments (compare with **F**).

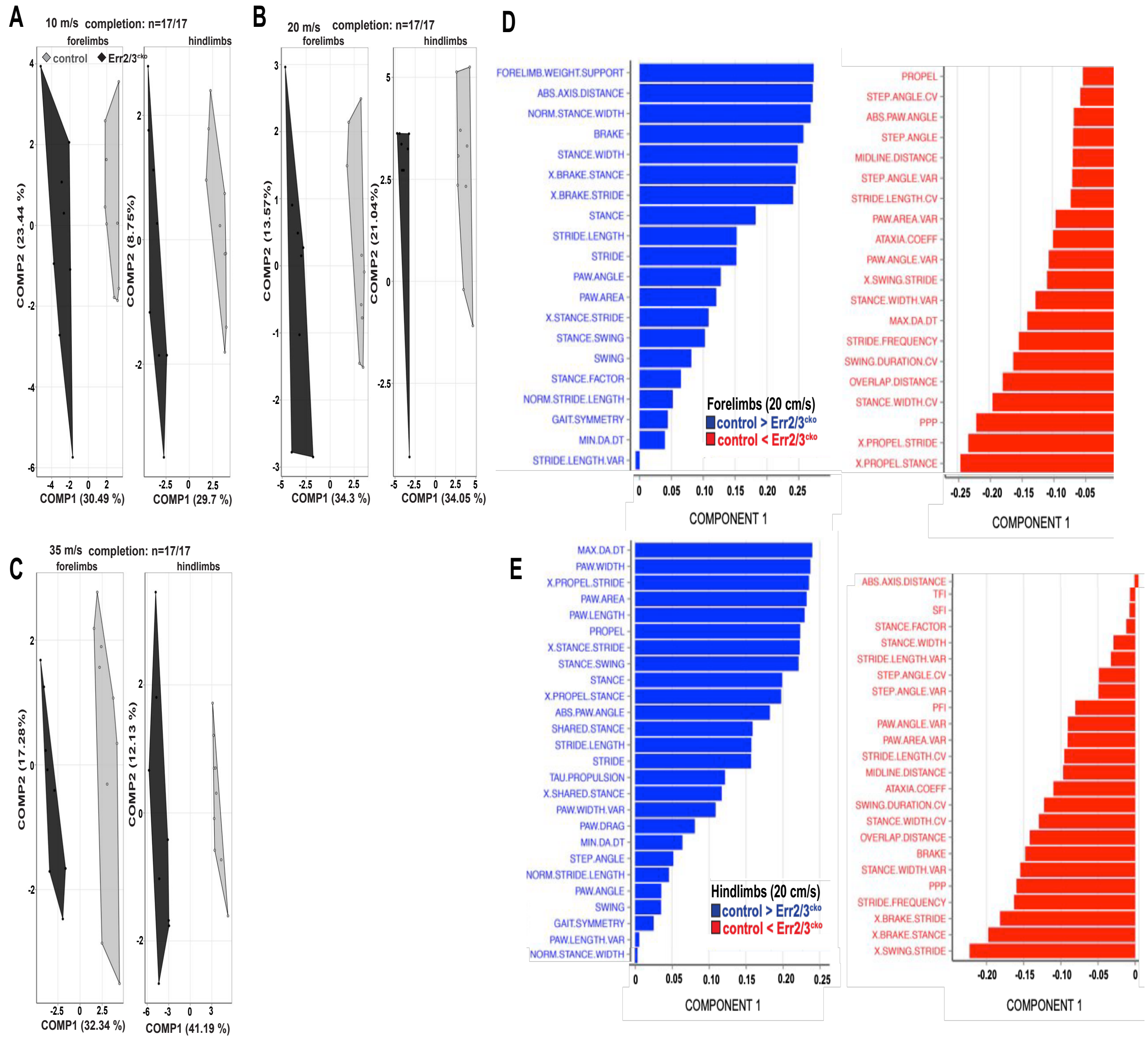


**Figure S8**. Gait alterations in Err2/3$^{cko}$ mice.

**(A-C)** Polygon graphs based on partial least squares (PLS) analysis of 58 gait variables measured during treadmill locomotion at 10 m•s$^{-1}$ **(A)**, 20 m•s$^{-1}$ **(B)** and 35 m•s$^{-1}$ **(C)**. Optimized model prediction was used to assign data sets for fore and hind limbs to either genotype (control versus Err2/3$^{cko}$) and the two components of the models were plotted against each other. Each one of the tested animals is represented by a single dot, while polygons group the animals of the same genotype. The amount of between-groups variance explained by each component in the model is expressed in percent of the total between-groups variance. The two components of our optimized models captured more than 25% of the variance in the predictors in both fore and hind limb at all treadmill speeds. These scores indicate that the method was able to capture the maximum variance between genotypes in the first dimension, which is also shown by the absence of overlap between the two groups on the x-axis, together indicating that Err2/3$^{cko}$ exhibit significant gait alterations compared to control mice. Yet, all Err2/3$^{cko}$ mice analyzed were able to successfully complete the treadmill locomotion tasks at all speeds tested (provided as the number of animals "n" running until "completion" for each speed). **(D, E)** Ranking of the variables' predictive capacities in the forelimb **(D)** and hindlimb **(E)** models. The most predictive parameters display the highest loadings (arbitrary units) independent of their sign. The mean value of all mice from the same group was calculated for each parameter and depicted in the bar charts. The sign of the loadings only indicates the direction in which gait parameters are affected (increased, red, or reduced, blue, in $Err2/3^{cko}$ compared to control mice).

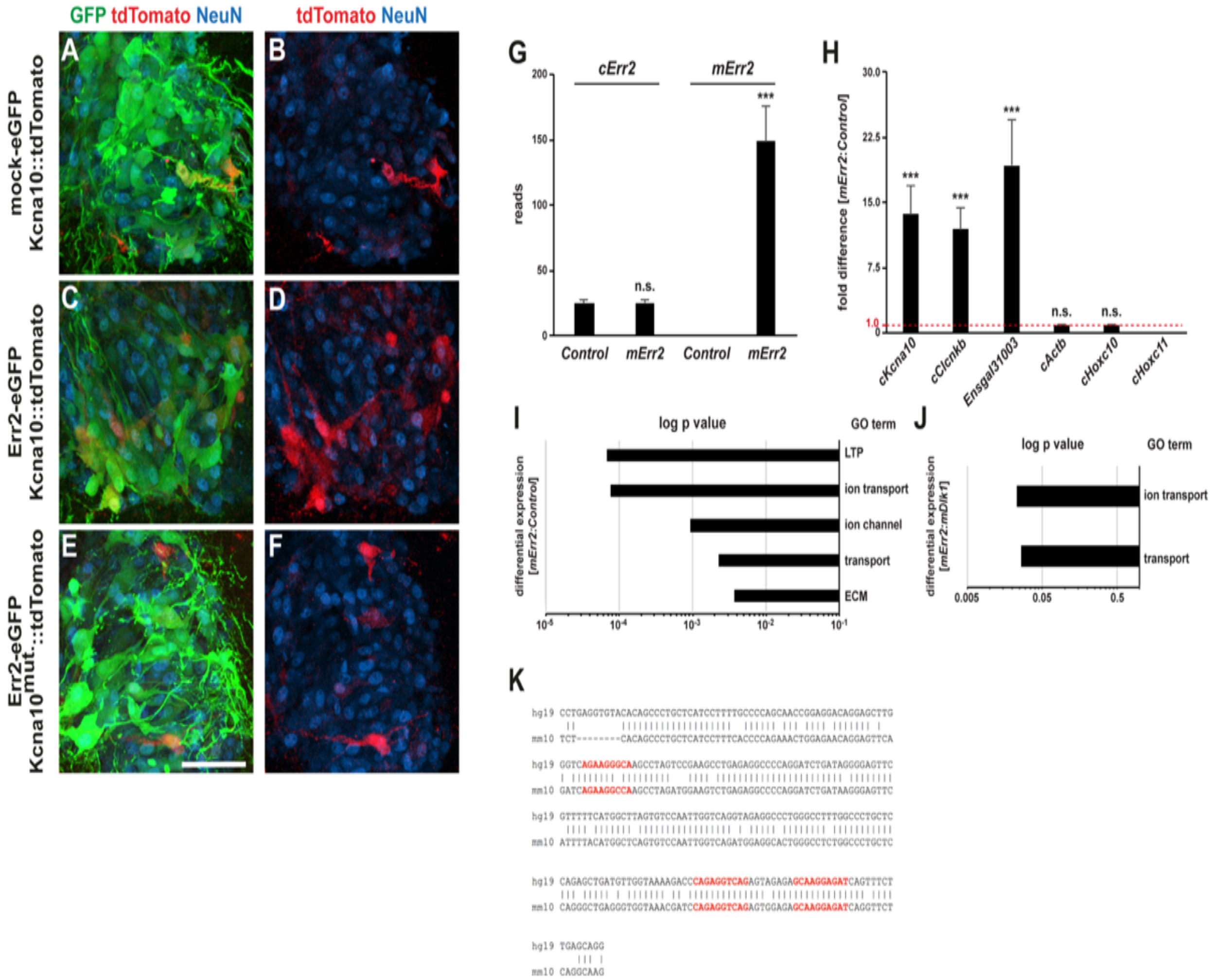


**Figure S9**. Err2 promotes a gene expression signature comprising determinants of membrane electrophysiological properties in chick motor neurons, including direct activation of *Kcna10*.

**(A-F)** Examples of tdTomato expression driven from a minimal promoter-tdTomato expression construct by the *Kcna10* ECR upon co-transfecting eGFP only ("mock") (control) **(C, D)**, Err2 and eGFP **(E, F)**, as well as the *Kcna10* ECR with scrambled Err2/3 binding sites plus Err2 and eGFP **(G, H)** (scale bar: 50 µm). **(G)** Numbers of RNA reads for endogenous chick *Err2* (*cErr2*) and mouse *Err2* (*mErr2*) in control (eGFP only) expressing chick motor neurons or chick motor neurons expressing mouse *Err2* (*mErr2*): RNA sequencing reveals unaltered cErr2 expression by forced *mErr2* expression, no reads of *mErr2* in the control sample (as expected) and ~6-fold over-expression of *mErr2* relative to endogenous *cErr2* levels. **(H)** Examples of genes upregulated by *mErr2* in chick motor neurons (given in fold change over control motor neurons), including genes encoding the voltage-gated potassium and chloride channels *Kcna10* and *cClcnkb*, respectively, as well as the not yet annotated gene *Ensgal00000031003*. Expression of genes related to motor pool identities, such as *Hoxa11* and *Hoxc11*, did not significantly change. **(I)** Clustered David functional annotations of genes differentially regulated by forced *mErr2* expression in chick motor neurons ("clustered" refers to genes associated with more than one annotated GO term), significance of classification is given as logarithmic p value. Gene ontology terms include "LTP" ("long-term potentiation": glutamate receptor *Grin2a*, neurotrophin receptor *Ntrk2*, sodium/potassium/calcium exchanger *Slc24a2* and tenascin R-encoding *Tnr*), "ion transport" and "ion channel" (*cClcnkb*, *Kcna10*, *Grin2a*, *Slc24a2,* potassium channel tetramerization domain containing *Kctd4*, gamma-aminobutyric acid receptor subunit *Gabrd*, voltage-gated sodium channel subunit *Scn5a*, voltage-gated potassium channel *Kcnh1*, transferrin, *TF* and Tweety-homolog 3-encoding *Ttyh3*), "transport" (adding translocase of inner mitochondrial membrane 10 homolog *TIMM10*, mitochondrial adenine nucleotide translocator *SLC25A4* to the "ion transport" list) and extracellular matrix (*Tnr*, Netrin1 *Ntn1* and aggrecan-encoding *Acan*). **(J)** GO terms for the genes differentially regulated by Err2 after subtracting the genes differentially regulated by Dlk1, leaving "ion transport" and "transport" as the only significantly clustered GO terms (including *Kcna10*, *cClcnkb*, *TF*, *Kctd4*). Note: while many if not all of these genes may in net contribute to the electrophysiological properties promoted by Err2 in motor neurons for practical reasons we focused on the single target gene *Kcna10* after finding that it recapitulated many of the effects of Err3 on motor neuron electrophysiological properties. **(K)** Alignment of evolutionary conserved on-coding genomic region (ECR) upstream of the *Kcna10* transcription start site between human and mouse. Predicted Err2/3 binding sites are highlighted in red (see Figure 7B).

**Supplementary Tables S1**

Summary of parameters recorded via whole-cell patch-clamping of mouse motor neurons in acute spinal cord slices. Genotypes of the recorded animals are given in the table.

| Membrane Property | control $FG^{high}$ (n=24) | control $FG^{low}$ (n=22) | $Err2/3^{cko}$ $FG^{high}$ (n=18) | $Err2/3^{cko}$ $FG^{low}$ (n=9) |
|---|---|---|---|---|
| Mean input resistance [MΩ] | 136.62 ± 14.63 | 36.55 ± 4.16 $^{a}$ *** | 114.48 ± 17.89 $^{b}$ n.s. | 35.79 ± 7.62 $^{c}$ n.s. |
| Mean capacitance [pF] | 76.07 ± 6.01 | 239.1 ± 17.17 $^{a}$ *** | 117.67 ± 8.93 $^{b}$ *** | 247.85 ± 22.19 $^{c}$ n.s. |
| Mean rheobase [pA] | 221.87 ± 31.34 | 909.09 ± 82.11 $^{a}$ *** | 419.72 ± 84.11 $^{b}$ * | 855.55 ± 124.85 $^{c}$ n.s. |
| Mean AHP amplitude [mV] | 3.23 ± 0.45 | 4.57 ± 0.28 $^{a}$ * | 3.69 ± 0.56 $^{b}$ n.s. | 4.27 ± 0.30 $^{c}$ n.s. |
| Mean AHP half-width [ms] | 32.22 ± 2.36 | 39.76 ± 2.75 $^{a}$ * | 45.31 ± 3.03 $^{b}$ *** | 33.55 ± 3.27 $^{c}$ n.s. |
| Mean AHP half-decay [ms] | 22.10 ± 1.89 | 27.78 ± 1.94 $^{a}$ * | 30.08 ± 2.50 $^{b}$ * | 23.89 ± 2.55 $^{c}$ n.s. |
| Mean firing frequency [Hz] | 50.65 ± 3.23 | 20.41 ± 1.70 $^{a}$ *** | 27.61 ± 3.15 $^{b}$ *** | 25.03 ± 3.76 $^{c}$ n.s. |
| Mean steady-state firing frequency [Hz] | 49.37 ± 3.20 | 20.18 ± 1.32 $^{a}$ *** | 26.25 ± 3.10 $^{b}$ *** | 24.99 ± 2.42 $^{c}$ n.s. |
| Mean instantaneous firing frequency [Hz] | 104.88 ± 6.90 | 30.09 ± 3.79 $^{a}$ *** | 56.52 ± 7.86 $^{b}$ *** | 35.83 ± 5.90 $^{c}$ n.s. |
| Mean gain [Hz/nA] | 161.01 ± 9.77 | 32.64 ± 4.02 $^{a}$ *** | 84.01 ± 12.34 $^{b}$ *** | 43.04 ± 9.81 $^{c}$ n.s. |
| Mean steady-state gain [Hz/nA] | 155.12 ± 9.87 | 31.66 ± 3.57 $^{a}$ *** | 78.01 ± 12.25 $^{b}$ *** | 41.86 ± 8.81 $^{c}$ n.s. |
| Mean instantaneous gain [Hz/nA] | 350.57 ± 17.15 | 70.62 ± 15.22 $^{a}$ *** | 215.45 ± 22.58 $^{b}$ *** | 79.15 ± 23.97 $^{c}$ n.s. |

Values show mean ± standard error of the mean (S.E.M.).

$^{a}$ indicates significant difference between control $FG^{high}$ (n=24) and control $FG^{low}$ (n=22) (Student's t-test);

$^{b}$ indicates significant difference between control $FG^{high}$ (n=24) and $Err2/3^{cko}$ $FG^{high}$ (n=18) (Student's t-test);

$^{c}$ indicates significant difference between control $FG^{low}$ (n=22) and $Err2/3^{cko}$ $FG^{low}$ (n=9) (Student's t-test);

***p-value <0.001; **p-value <0.01; *p-value <0.05; n.s., not significant. n= # of neurons.

**Supplementary Table S2**

Summary of parameters recorded via whole-cell patch-clamping of chick motor neurons in acute spinal cord slices. Constructs used to stably transfect the motor neurons prior to the recordings are in the table.

| Membrane Property | CMV-eGFP-control 1 (n=21) | CMV-VP16-Err2-eGFP (n=20) | CMV-Err2-eGFP (n=21) | CMV-Err3-eGFP (n=21) | CMV-EnR-Err2-eGFP (n=17) | CMV-eGFP-control 2 (n=23) | CMV-KCNA10-eGFP (n=24) |
|---|---|---|---|---|---|---|---|
| Mean input resistance [MΩ] | 272.13 ± 34.62 | 298.72 ± 26.67 [a] n.s. | 309.96 ± 28.96 [a] n.s. | 299.52 ± 31.54 [a] n.s. | 181.48 ± 14.14 [a] * | 263.26 ± 31.59 [b] n.s. | 373.99 ± 47.38 [c] n.s. |
| Mean capacitance [pF] | 237.77 ± 14.82 | 141.63 ± 7.38 [a] *** | 185.20 ± 7.96 [a] *** | 189.54 ± 7.33 [a] *** | 218.26 ± 8.91 [a] n.s. | 241.29 ± 12.01 [b] n.s. | 183.96 ± 9.74 [c] *** |
| Mean rheobase [pA] | 160.95 ± 25.85 | 61 ± 6.36 [a] *** | 114.28 ± 15.10 [a] n.s. | 128.28 ± 23.41 [a] n.s. | 182.94 ± 18.81 [a] n.s. | 160.43 ± 19.97 [b] n.s. | 67.91 ± 11.15 [c] *** |
| Mean AHP amplitude [mV] | 8.48 ± 0.75 | 5.85 ± 0.51 [a] *** | 6.54 ± 0.81 [a] n.s. | 5.44 ± 0.51 [a] *** | 8.17 ± 0.61 [a] n.s. | 7.52 ± 0.58 [b] n.s. | 6.33 ± 0.54 [c] n.s. |
| Mean AHP half-width [ms] | 118.53 ± 9.37 | 97.27 ± 5.57 [a] n.s. | 110.39 ± 7.13 [a] n.s. | 110.62 ± 4.01 [a] n.s. | 94.80 ± 4.06 [a] *** | 130.96 ± 8.22 [b] n.s. | 121.55 ± 8.14 [c] n.s. |
| Mean AHP half-decay [ms] | 85.97 ± 7.16 | 69.91 ± 4.26 [a] n.s. | 75.60 ± 5.54 [a] n.s. | 77.0 ± 3.22 [a] n.s. | 67.49 ± 3.12 [a] * | 93.34 ± 5.88 [b] n.s. | 87.34 ± 6.17 [c] n.s. |
| Mean firing frequency [Hz] | 13.63 ± 0.63 | 19.82 ± 1.10 [a] *** | 15.91± 0.78 [a] * | 16.93 ± 1.12 [a] *** | 15.08 ± 0.87 [a] n.s. | 12.69 ± 0.73 [b] n.s. | 15.83 ± 0.57 [c] *** |
| Mean steady-state firing frequency [Hz] | 13.26 ± 0.62 | 19.14 ± 1.06 [a] *** | 15.66 ± 1.05 [a] * | 16.65 ± 0.67 [a] *** | 14.42 ± 0.87 [a] n.s. | 12.51 ± 0.73 [b] n.s. | 15.49 ± 0.56 [c] *** |
| Mean instantaneous firing frequency [Hz] | 14.87 ± 1.05 | 32.01 ± 2.36 [a] *** | 18.39 ± 1.42 [a] * | 22.0 ± 1.26 [a] *** | 14.79 ± 1.36 [a] n.s. | 14.12 ± 1.22 [b] n.s. | 21.46 ± 0.94 [c] *** |
| Mean gain [Hz/nA] | 44.84 ± 2.75 | 101.01 ± 6.58 [a] *** | 58.0 ± 3.49 [a] *** | 62.47 ± 2.66 [a] *** | 44.58 ± 5.14 [a] n.s. | 42.45 ± 2.63 [b] n.s. | 46.57 ± 2.69 [c] n.s. |
| Mean stead-state gain [Hz/nA] | 44.35 ± 2.79 | 93.21 ± 6.13 [a] *** | 57.12 ± 4.33 [a] *** | 60.75 ± 2.69 [a] *** | 43.99 ± 5.43 [a] n.s. | 42.1 ± 2.63 [b] n.s. | 44.26 ± 2.76 [c] n.s. |
| Mean instantaneous gain [Hz/nA] | 60.86 ± 5.67 | 209.59 ± 16.68 [a] *** | 79.59± 8.88 [a] n.s. | 106.75 ± 8.27 [a] *** | 54.67± 8.99 [a] n.s. | 58.07 ± 5.20 [b] n.s. | 85.72 ± 5.36 [c] *** |

CMV-eGFP-control 1 versus CMV-VP16-Err2-eGFP, CMV-Err2-eGFP, CMV-Err3-eGFP, CMV-EnR-Err2-eGFP and CMV-eGFP-control 2 versus CMV-KCNA10-eGFP.

Values show mean ± standard error of the mean (S.E.M.).

[a] indicates significant difference compared to CMV-eGFP-control 1 (Student's t-test);

[b] indicates significant difference between CMV-eGFP-control 1 and CMV-eGFP-control 2 (Student's t-test);

[c] indicates significant difference between CMV-eGFP-control 2 and CMV-KCNA10-eGFP (Student's t-test); ***p-value <0.001; **p-value <0.01; *p-value <0.05; n.s., not significant.

**Supplementary Table S3: Gene expression signatures promoted by forced Err2:Vp16 expression**

**Err2 vs. Control: Differentially Regulated (adj. P value <0.05)**

| ENSEMBL_ID | GENE | baseMean_CMV-eGFP | baseMean_CMV-VP16-Err2 | log2FoldChange | padj |
|---|---|---|---|---|---|
| mERR2 | NA | 0 | 148,8927158 | 2,469843079 | 8,72E-43 |
| ENSGALG00000033638 | SLC26A7 | 4,265182372 | 50,56759488 | 1,317066411 | 4,82E-11 |
| ENSGALG00000031003 | NA/lincRNA | 12,57733457 | 241,5743348 | 0,878204649 | 5,00E-05 |
| ENSGALG00000026687 | PLCD4 | 30,86611927 | 79,76434748 | 0,868225991 | 0,000266119 |
| ENSGALG00000000441 | KCNA10 | 1,672112917 | 22,83046587 | 0,840306626 | 0,000233827 |
| ENSGALG00000003713 | CLCNKB | 1,374411237 | 21,44579174 | 0,750353053 | 0,001014112 |
| ENSGALG00000006453 | TF | 363,0028699 | 1277,800233 | 0,743620697 | 0,009595313 |
| ENSGALG00000030349 | NA/lincRNA | 21,59581471 | 45,04577184 | 0,69172147 | 0,016732209 |
| ENSGALG00000035927 | ST8SIA5 | 118,3581207 | 210,8200842 | 0,691238907 | 0,000258701 |
| ENSGALG00000023626 | NTN1 | 144,415863 | 321,5220373 | 0,65927369 | 0,039598485 |
| ENSGALG00000032650 | NA/lincRNA | 2,738608622 | 46,37068414 | 0,62468058 | 0,009595313 |
| ENSGALG00000038087 | CACNG3 | 265,3702433 | 437,379539 | 0,623646543 | 0,016600396 |
| ENSGALG00000016567 | EPHX2 | 88,77353148 | 158,2518529 | 0,620198457 | 0,020735307 |
| ENSGALG00000037586 | YIF1B | 76,68732207 | 142,1448594 | 0,608826972 | 0,041912825 |
| ENSGALG00000037586 | NA | 76,68732207 | 142,1448594 | 0,608826972 | 0,041912825 |
| ENSGALG00000021552 | RASL10A | 1,349370367 | 8,977138233 | 0,593304486 | 0,038691724 |
| ENSGALG00000021552 | RASL10A | 1,349370367 | 8,977138233 | 0,593304486 | 0,038691724 |
| ENSGALG00000031333 | NA/lincRNA | 54,05032414 | 89,55651463 | 0,564248969 | 0,038691724 |
| ENSGALG00000003904 | MLPH | 57,28605637 | 94,38629697 | 0,562638157 | 0,026621491 |
| ENSGALG00000042809 | NDUFV1 | 463,9242548 | 734,8015889 | 0,534525008 | 0,034569352 |
| ENSGALG00000042809 | NDUFV1 | 463,9242548 | 734,8015889 | 0,534525008 | 0,034569352 |
| ENSGALG00000026518 | RUNDC3A | 388,5725554 | 591,1973078 | 0,50029069 | 0,039598485 |
| ENSGALG00000037402 | CTGF | 254,0126666 | 360,9906462 | 0,433903534 | 0,046425712 |
| ENSGALG00000043661 | NA | 244,1591247 | 179,8885243 | -0,385097238 | 0,034569352 |
| ENSGALG00000043661 | NA | 244,1591247 | 179,8885243 | -0,385097238 | 0,034569352 |
| ENSGALG00000014337 | NWD2 | 1120,190542 | 804,5079487 | -0,423698149 | 0,039598485 |
| ENSGALG00000015080 | SLC24A2 | 180,1718787 | 123,1711858 | -0,485086299 | 0,044784742 |
| ENSGALG00000002555 | RET | 1988,865618 | 1303,385823 | -0,514322756 | 0,020735307 |
| ENSGALG00000009877 | KCNH1 | 139,7928925 | 90,49469093 | -0,56953159 | 0,017372822 |
| ENSGALG00000015953 | NA | 122,1004061 | 73,47563433 | -0,579184828 | 0,022890452 |
| ENSGALG00000011860 | NA | 155,6867278 | 91,88746368 | -0,594016264 | 0,009595313 |
| ENSGALG00000038225 | SEMA3E | 209,9067866 | 127,5941239 | -0,596481046 | 0,002300999 |
| ENSGALG00000007647 | MYO3A | 276,01235 | 161,2458995 | -0,612768832 | 0,009595313 |
| ENSGALG00000007666 | KCHIP2 | 272,4037453 | 153,3790909 | -0,620080381 | 0,020735307 |
| ENSGALG00000011235 | GALNT15 | 157,4091888 | 83,87246896 | -0,641352356 | 0,022537939 |
| ENSGALG00000005122 | MYO1H | 37,90794484 | 11,06875412 | -0,686774368 | 0,022890452 |
| ENSGALG00000007079 | PAPPA | 466,5121276 | 215,4060778 | -0,689555327 | 0,020735307 |
| ENSGALG00000004526 | TNR | 1302,722899 | 725,484286 | -0,700084494 | 0,014265458 |
| ENSGALG00000016804 | SLC5A7 | 933,3251133 | 392,2052743 | -0,704256721 | 0,020735307 |
| ENSGALG00000006112 | SCN5A | 228,7238968 | 129,0247733 | -0,714573054 | 0,004621824 |
| ENSGALG00000007278 | GRIN2A | 376,5548026 | 187,2086824 | -0,726664413 | 0,039712878 |

**Err2 vs. Dlk1: Differentially Regulated (adj. P value <0.05)**

| ENSEMBL_ID | GENE | Mean_Exp2-pCaggs-Mean_Exp2-CMV-VP1 | | log2FoldChange | padj |
|---|---|---|---|---|---|
| mERR2 | NA | 0,347906688 | 111,4544545 | 1,89621077 | 9,05E-36 |
| ENSGALG00000006453 | TF | 314,3396102 | 1232,834664 | 0,895184392 | 3,91E-06 |
| ENSGALG00000033638 | SLC26A7 | 4,560647586 | 38,87943958 | 0,782518954 | 7,74E-06 |
| ENSGALG00000026687 | PLCD4 | 25,47441187 | 61,67822153 | 0,670239059 | 0,003299043 |
| ENSGALG00000035927 | ST8SIA5 | 92,5502537 | 162,4795051 | 0,651534365 | 0,000146488 |
| ENSGALG00000034741 | ETNPPL | 15,41151319 | 38,03236147 | 0,618074707 | 0,012177036 |
| ENSGALG00000003713 | NA | 1,270882488 | 17,62442709 | 0,590832101 | 0,000506105 |
| ENSGALG00000010116 | DNAH8 | 14,55863737 | 29,43089254 | 0,581380768 | 0,023532367 |
| ENSGALG00000010614 | SLC25A4 | 323,2368851 | 586,0961243 | 0,536563778 | 0,042032009 |
| ENSGALG00000041255 | ADAM12 | 78,83611166 | 138,0966622 | 0,533139892 | 0,038494488 |
| ENSGALG00000031003 | NA | 14,22690137 | 220,9880052 | 0,531922859 | 0,001462047 |
| ENSGALG00000005046 | SHMT1 | 61,13744112 | 108,365422 | 0,528420289 | 0,041528512 |
| ENSGALG00000043614 | NA | 36,72089105 | 59,82975244 | 0,498297097 | 0,04900555 |
| ENSGALG00000012361 | NIN | 224,5431866 | 337,6879396 | 0,480420629 | 0,014297375 |
| ENSGALG00000032650 | NA | 0,91622273 | 30,52935914 | 0,464417297 | 0,002738721 |
| ENSGALG00000000441 | KCNA10 | 2,032050564 | 18,51764808 | 0,426827634 | 0,021987807 |
| ENSGALG00000013865 | IFNGR1 | 159,7391982 | 225,1183169 | 0,419255147 | 0,043015604 |
| ENSGALG00000002056 | TANGO2 | 477,3940829 | 639,1150303 | 0,385271357 | 0,003299043 |
| ENSGALG00000042519 | NA | 219,4340852 | 294,5128228 | 0,382016811 | 0,017815937 |
| ENSGALG00000026364 | ASAH1 | 864,5273052 | 1176,704164 | 0,38031213 | 0,044461297 |
| ENSGALG00000010163 | LGR5 | 216,9245197 | 283,2494123 | 0,356888149 | 0,033857654 |
| ENSGALG00000010681 | CPT2 | 458,5181001 | 599,2033981 | 0,345220807 | 0,027849386 |
| ENSGALG00000002216 | NEK7 | 471,6497678 | 591,868455 | 0,298647144 | 0,028292802 |
| ENSGALG00000031034 | NA | 611,3165617 | 725,1362468 | 0,236429862 | 0,028471659 |
| ENSGALG00000005237 | FGD3 | 2232,502657 | 1914,209806 | -0,213217196 | 0,034167133 |
| ENSGALG00000041621 | LY6E | 1708,696881 | 1363,715726 | -0,302318527 | 0,041528512 |
| ENSGALG00000026091 | CACNG4 | 543,3716677 | 429,817395 | -0,308900307 | 0,003421032 |
| ENSGALG00000014485 | LDB2 | 425,0461372 | 335,3542869 | -0,311476887 | 0,010062561 |
| ENSGALG00000012823 | TRIM24 | 738,3253272 | 582,3390809 | -0,314802855 | 0,002738721 |
| ENSGALG00000035065 | FAM110B | 596,2350028 | 471,1176527 | -0,321575836 | 0,0064062 |
| ENSGALG00000029011 | SLC35F1 | 799,9661685 | 617,1298904 | -0,336175255 | 0,029236493 |
| ENSGALG00000040131 | CAMKK2 | 751,0564962 | 580,2799522 | -0,336667142 | 0,021213308 |
| ENSGALG00000043764 | KCNS2 | 465,7203701 | 359,0391666 | -0,339502057 | 0,015599686 |
| ENSGALG00000010470 | TCERG1L | 344,8327809 | 261,0919878 | -0,347157404 | 0,027849386 |
| ENSGALG00000009499 | NPY5R | 214,5188726 | 163,3418759 | -0,357863884 | 0,046195415 |
| ENSGALG00000029510 | STK32B | 412,1111364 | 312,0357743 | -0,369631847 | 0,002738721 |
| ENSGALG00000030481 | RAB40C | 205,205237 | 154,0809613 | -0,374581373 | 0,006807235 |
| ENSGALG00000011721 | AKAP5 | 362,6730819 | 269,2107832 | -0,378770216 | 0,003561195 |
| ENSGALG00000032418 | RAB26 | 891,7519309 | 664,6439139 | -0,378915383 | 0,024299634 |
| ENSGALG00000039145 | CA8 | 207,9260445 | 151,1368697 | -0,39208299 | 0,028292802 |
| ENSGALG00000028682 | CBLN2 | 2700,681897 | 1977,470378 | -0,406038319 | 0,003236376 |
| ENSGALG00000005319 | CDH8 | 1590,599778 | 1143,90989 | -0,406639041 | 0,022379658 |
| ENSGALG00000028602 | NA | 1678,683179 | 1243,913626 | -0,406685337 | 3,79E-05 |
| ENSGALG00000014930 | ENC1 | 1415,348845 | 996,1477587 | -0,415004797 | 0,046195415 |
| ENSGALG00000030607 | CHST11 | 179,0617537 | 124,1911405 | -0,417856341 | 0,042164828 |
| ENSGALG00000039595 | BTBD11 | 448,0902106 | 314,7504676 | -0,442154143 | 0,003299043 |
| ENSGALG00000043240 | ARPP21 | 2555,320231 | 1714,573359 | -0,449572525 | 0,047090096 |
| ENSGALG00000031058 | NA | 90,11008863 | 62,92204581 | -0,454851488 | 0,02275113 |
| ENSGALG00000016975 | KCTD4 | 169,7271499 | 114,2735207 | -0,458431362 | 0,037191435 |
| ENSGALG00000009556 | PRICKLE1 | 427,3845665 | 292,5713101 | -0,458653812 | 0,006333455 |
| ENSGALG00000011244 | DLK1 | 249,3847488 | 165,1331459 | -0,4687905 | 0,027849386 |
| ENSGALG00000009738 | NA | 244,3651616 | 162,8039211 | -0,473406574 | 0,016526823 |
| ENSGALG00000009021 | ST6GALNAC5 | 439,5680912 | 296,8923737 | -0,482507986 | 0,002738721 |
| ENSGALG00000026167 | PIK3R5 | 424,1382713 | 272,4541839 | -0,51473103 | 0,006333455 |
| ENSGALG00000005657 | CRHR2 | 180,1732623 | 118,7708259 | -0,516956452 | 0,000361141 |
| ENSGALG00000040675 | HS3ST2 | 150,301388 | 86,94919033 | -0,526278494 | 0,039080913 |
| ENSGALG00000032554 | C1QL3 | 33,96109487 | 17,90109227 | -0,538792077 | 0,046043563 |
| ENSGALG00000012319 | NA | 235,5338175 | 146,0376512 | -0,538889051 | 0,003760199 |
| ENSGALG00000029557 | NA | 60,74993739 | 30,70228324 | -0,539689873 | 0,046108833 |

| | | | | | |
|---|---|---|---|---|---|
| ENSGALG00000040582 | LMX1B | 288,2401077 | 136,9632154 | -0,543005604 | 0,04900555 |
| ENSGALG00000003699 | EBF1 | 152,1117135 | 77,28746611 | -0,545717984 | 0,042164828 |
| ENSGALG00000015419 | PENK | 1036,71436 | 632,383155 | -0,545988667 | 0,006333455 |
| ENSGALG00000030822 | DRGX | 192,1930168 | 63,16113094 | -0,547938546 | 0,032283554 |
| ENSGALG00000025950 | PRKCA | 3651,41106 | 2015,55591 | -0,569001309 | 0,016526823 |
| ENSGALG00000012542 | RASD2 | 94,86097218 | 37,63268903 | -0,590212147 | 0,021809011 |
| ENSGALG00000012481 | TNFAIP6 | 47,76153114 | 23,03690277 | -0,599503893 | 0,016090026 |
| ENSGALG00000035058 | CAMKV | 416,9321466 | 231,833726 | -0,610330039 | 0,003299043 |
| ENSGALG00000009737 | tac1 | 1443,622928 | 790,5767616 | -0,614840569 | 0,003236376 |
| ENSGALG00000017411 | NA | 48,56308477 | 23,72855669 | -0,665404616 | 0,002120509 |
| ENSGALG00000021589 | RTN4RL1 | 118,3672524 | 60,88109905 | -0,669766575 | 0,001191387 |
| ENSGALG00000033412 | NA | 522,2640794 | 250,7248322 | -0,68161486 | 0,001539255 |
| ENSGALG00000003034 | SS2 | 386,1792747 | 160,1397805 | -0,682712431 | 0,002742908 |
| ENSGALG00000026264 | MAFA | 437,7994622 | 177,7669082 | -0,694985932 | 0,002402827 |
| ENSGALG00000013604 | VIP | 142,3794886 | 67,2003381 | -0,768032835 | 1,85E-05 |
| mDLK1 | NA | 581,1854666 | 0,256510976 | -2,406946313 | 1,25E-55 |

**Supplementary Movies S1-S4**

**Supplementary Movie S1**

Example of a control mouse navigating a horizontal beam. The animal movement was tracked using GoPro HD Hero2 (GoPro Inc., San Mateo U.S.A) fitted to a custom-built slider track. The video's frame rate (acquired at 120 fps) was reduced to 60 fps and speed was slowed to 25-40% of the original speed.

**Supplementary Movie S2**

Example of a *Err2/3*$^{cko}$ mouse navigating a horizontal beam. The video was acquired and modified as above (*Supplementary Movie S1*).

**Supplementary Movie S3**

Example of a control mouse navigating a horizontal ladder. The video was acquired and modified as above (*Supplementary Movie S1*).

**Supplementary Movie S4**

Example of a *Err2/3*$^{cko}$ mouse navigating a horizontal ladder. The video was acquired and modified as above (*Supplementary Movie S1*).
